# Broad therapeutic benefit of myosin inhibition in hypertrophic cardiomyopathy

**DOI:** 10.1101/2024.03.22.584986

**Authors:** Laura Sen-Martín, Ángel Fernández-Trasancos, Miguel Á. López-Unzu, Divya Pathak, Alessia Ferrarini, Verónica Labrador-Cantarero, David Sánchez-Ortiz, María Rosaria Pricolo, Natalia Vicente, Diana Velázquez-Carreras, Lucía Sánchez-García, Jose Ángel Nicolás-Ávila, María Sánchez-Díaz, Saskia Schlossarek, Lorena Cussó, Manuel Desco, María Villalba-Orero, Gabriela Guzmán-Martínez, Enrique Calvo, Roberto Barriales-Villa, Jesús Vázquez, Fátima Sánchez-Cabo, Andrés Hidalgo, Lucie Carrier, James A. Spudich, Kathleen M. Ruppel, Jorge Alegre-Cebollada

**Affiliations:** Centro Nacional de Investigaciones Cardiovasculares Carlos III (CNIC), Madrid, Spain; Department of Biochemistry and Stanford Cardiovascular Institute, Stanford University School of Medicine; Servicio de Cardiología, Hospital Puerta de Hierro de Majadahonda, Madrid, Spain; Department of Experimental Pharmacology and Toxicology, University Medical Center Hamburg-Eppendorf, Hamburg, Germany; 2DZHK (German Centre for Cardiovascular Research), partner site Hamburg/Kiel/Lübeck, Hamburg, Germany; Laboratorio de imagen para pequeño animal de experimentación, Instituto de Investigación Sanitaria Gregorio Marañón, Madrid, Spain; CIBER de salud mental, Instituto de salud Carlos III, Madrid, Spain; Departamento de Bioingeniería, Universidad Carlos III de Madrid, Madrid, Spain; Departamento de Medicina y Cirugía Animal, Facultad de Veterinaria, Universidad Complutense de Madrid, Spain; Department of Cardiology, La Paz University Hospital, IdiPaz, Madrid, Spain; Department of Medicine, Faculty of Biomedical and Health Sciences, Universidad Europea de Madrid, Spain; Unidad de Cardiopatías Familiares, Servicio de Cardiología, Complexo Hospitalario Universitario de A Coruña, Servizo Galego de Saúde (SERGAS); Instituto de Investigación Biomédica de A Coruña (INIBIC), Universidade da Coruña, A Coruña, Spain; CIBER de Enfermedades Cardiovasculares (CIBERCV), Madrid, Spain

## Abstract

Myosin inhibitor mavacamten is the only targeted treatment available for hypertrophic cardiomyopathy (HCM), a disease caused by hundreds of genetic variants that affect mainly sarcomeric myosin and its negative regulator cardiac myosin-binding protein C (cMyBP-C, encoded by *MYBPC3*). Here, we have examined whether the reported limited efficacy of mavacamten in a fraction of HCM patients can result from dissimilar HCM pathomechanisms triggered by different genetic variants, a scenario particularly relevant for *MYBPC3*-associated HCM. To this aim, we have generated knock-in mice including missense pathogenic variant cMyBP-C p.R502W, which, different from patients who carry truncations in the protein, develop progressive pathogenic myocardial remodeling in the absence of alterations of cMyBP-C levels and localization. Mechanistically, we find that mutation R502W reduces the binding affinity of cMyBP-C for myosin without inducing a shift towards more active myosin conformations as observed when cMyBP-C levels are reduced. Despite these diverging molecular alterations, we show that mavacamten blunts myocardial remodeling both in R502W and cMyBP-C-deficient, knock-out hearts. These beneficial effects are accompanied by improved tolerance to exercise only in R502W animals. Hence, our results indicate that myosin inhibition is effective to treat HCM caused by both truncating and missense variants in *MYBPC3* regardless of the primary pathomechanisms they elicit.

## INTRODUCTION

Hypertrophic cardiomyopathy (HCM) is the most frequent familial heart disease, affecting 0.5% of the population ^1,2^. HCM hearts show distinctive, but somewhat heterogeneous anatomical features ranging from myocardial crypts and hypertrabeculation to more common asymmetrical septal hypertrophy ^3^. At the histological level, HCM myocardium displays interstitial fibrosis and cardiomyocyte hypertrophy ^4^. Diastolic dysfunction is an early sign of the disease, while systolic alterations can occur in latter stages ^2^. Major risks of HCM include heart failure and sudden cardiac death ^2^. Despite the severity of these outcomes, clinical management of HCM has been traditionally limited to symptom alleviation, failing to prevent pathogenic myocardial remodeling ^5^.

HCM is inherited in an autosomal dominant manner typically implicating variants in proteins of the sarcomere, the contractile unit of cardiomyocytes. Most of the ∼40% HCM patients for which a genetic cause is identified carry variants in *MYH7* and *MYBPC3*, codifying respectively for the ATP-driven molecular motor β-myosin heavy chain ^6^, and cardiac myosin binding protein C (cMyBP-C), a modulator of myosin activity present in the C-zone of sarcomeres ^7,8^ (**Figure 1**). The partnership between these two proteins is further illustrated by the fact that cMyBP-C promotes myosin transition from the more active disordered relaxed (DRX) conformation to the super relaxed (SRX) state, which shows 10-fold slower ATPase activity ^9^.

**Figure 1.**
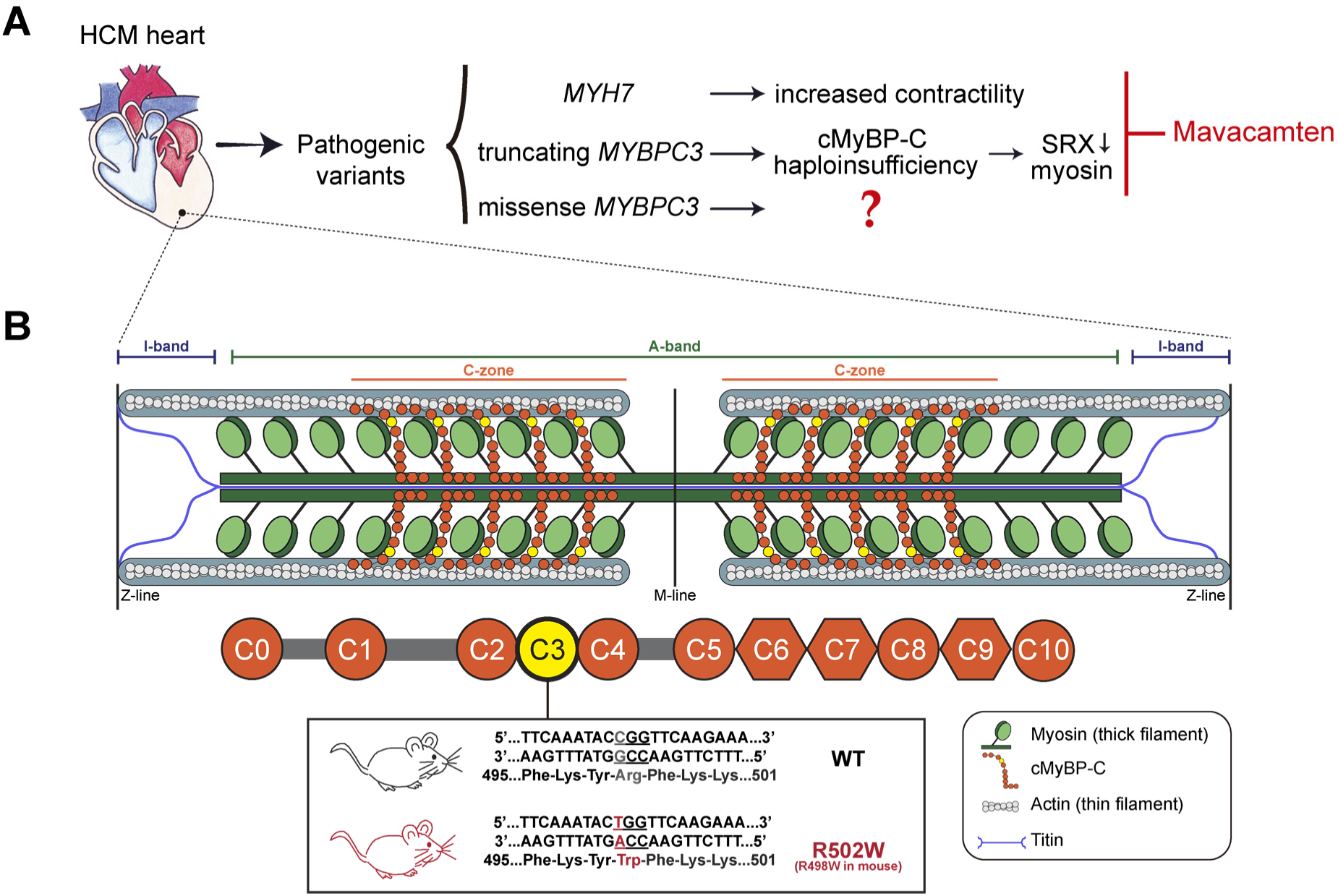
A mouse model of HCM induced by missense variant p.R502W in cMyBP-C. **(A)** Overview of the main types of pathogenic variants in *MYH7* and *MYBPC3* causing HCM. Variants in *MYH7* increase myosin contractility. Truncations in *MYBPC3* result in reduced cMyBP-C levels (haploinsufficiency) that lead to less myosin molecules in SRX conformation. The mechanisms by which missense variants in *MYBPC3* cause HCM remain poorly understood. The FDA-approved drug mavacamten inhibits myosin contractility and restores ratio of SRX/DRX myosin conformations. **(B)** Representation of the cardiac sarcomere highlighting its main regions and protein components. cMyBP-C, located at the C-zone, is a multidomain protein composed of 8 immunoglobulin-like and 3 fibronectin-like domains (represented as circles and hexagons, respectively). The human R502W variant targets the central C3 domain of the protein (colored in yellow) and is equivalent to variant R498W in mice.

The HCM myocardium is hypercontractile, particularly in the early stages of the disease, which raised the possibility that increased sarcomere activity is at the basis of HCM development ^2,5^. Indeed, pathogenic variants in *MYH7* have been shown to induce sarcomere hypercontractility through different mechanisms, such as boosting intrinsic myosin motor properties (contraction force, velocity, ATPase activity, proportion of DRX) and/or increasing the number of myosin heads available for interaction with actin ^6,10–13^ (**Figure 1A**). In the case of truncating variants in *MYBPC3*, cMyBP-C haploinsufficiency causes sarcomere hypercontractility *via* reduced proportion of SRX myosin ^14–17^ (**Figure 1A**). Whether missense *MYBPC3* variants also affect SRX/DRX myosin conformations, especially when cMyBP-C levels are preserved, remains unknown.

The concept that sarcomere hypercontractility underlies HCM progress has led to the development of therapies based on myosin inhibition ^18,19^. The success of this strategy has recently been demonstrated by the remarkable clinical benefits of myosin ATPase inhibitor mavacamten, the first-in-class of this new family of drugs, in the EXPLORER-HCM phase III clinical trial ^5,20^. In this study, patients with obstructive HCM showed clear benefits in a composite primary endpoint based on improved peak oxygen consumption and reduction in New York Heart Association functional class. However, more than 50% of patients in the mavacamten group failed to reach the primary endpoint of the study. The reasons for this heterogenous response remain unknown. A contributing factor could be the broad genetic landscape of patients, since the effects of myosin inhibition could vary between carriers of variants that induce different pathomechanisms ^21^. Indeed, considering that mavacamten efficacy has been linked to normalization of SRX/DRX myosin ratios ^16,18,22^, the possibility exists that clinical benefit may be limited in carriers of *MYBPC3* missense variants not compromising myosin SRX conformation (**Figure 1A**).

To examine this possibility, we first set out to generate a mouse model of HCM triggered by a *Mybpc3* variant that preserves cMyBP-C levels. We chose cMyBP-C p.R502W for several reasons. First, previous *in vitro* studies have shown that p.R502W does not result in changes in mRNA processing nor protein stability, two typical cMyBP-C haploinsufficiency drivers ^23,24^. In addition, p.R502W, the most common cause of HCM in humans found in up to 2% of cases ^25^, targets the central C3 domain of cMyBP-C (**Figure 1B**), a hotspot of similar, highly prevalent pathogenic *MYBPC3* variants ^26^. Using R502W mice, here we show that R502W hearts have preserved cMyBP-C levels and myosin SRX/DRX conformations but develop typical features of HCM, and demonstrate that mavacamten is able to blunt pathogenic myocardial remodeling in both cMyBP-C R502W and knock-out (KO) animals despite non-overlapping primordial HCM pathomechanisms.

## RESULTS

### HCM remodeling in homozygous R502W mice

The genomic region targeted by *MYBPC3* p.R502W (NM_000256.3:c.1504C>T) is highly conserved in human and mice (**Supplementary Figure S1A**). We used CRISPR-Cas9 to knock in the variant in murine *Mybpc3* (**Supplementary Figure S1B,C**). The resulting R502W mice breed normally and homozygous pups are obtained in Mendelian proportions (**Supplementary Figure S1D**). Considering that mice carrying truncating *MYBPC3* variants develop robust HCM remodeling only in homozygosis ^27^, we decided to focus on homozygous R502W animals. We found that explanted R502W hearts have a more rounded morphology than wild-type (WT) counterparts (**Figure 2A**). Indeed, *in vivo* echocardiography showed that R502W left ventricles have eccentricity indexes closer to 1 than WT at all examined time points (**Figure 2B,C**). Remarkably, R502W animals develop progressive cardiac hypertrophy, as determined both *in vivo* by echocardiography (**Supplementary Figure S2A**) and postmortem by measuring atria-free heart-weight (HW) to tibia length (TL) ratios (10-20% increase in HW/TL starting at 18 weeks of age, **Figure 2D**). Echocardiography also detected subtle interventricular septum and left ventricular posterior wall thickening in mutant mice (**Supplementary Figure S2B,C**). To characterize these anatomical changes at higher resolution, we used cardiac magnetic resonance imaging (MRI), which showed increased number of trabeculae in the left ventricle wall of R502W mice at 9, 18 and 80 weeks of age (**Figure 2E,F**), particularly in the apical, inferolateral and anterolateral regions (**Figure 2G**). At the histological level, we found that R502W cardiomyocytes have increased cross-sectional area at 18 weeks (**Figure 2H,I**), a phenotype that is more evident in 80-week-old mice (**Figure 2J,K**). Using Picrosirius red staining, we also detected increased interstitial deposition of collagen in R502W myocardium at both time points, indicative of cardiac fibrosis in mutant animals (**Figure 2L-O**).

**Figure 2.**
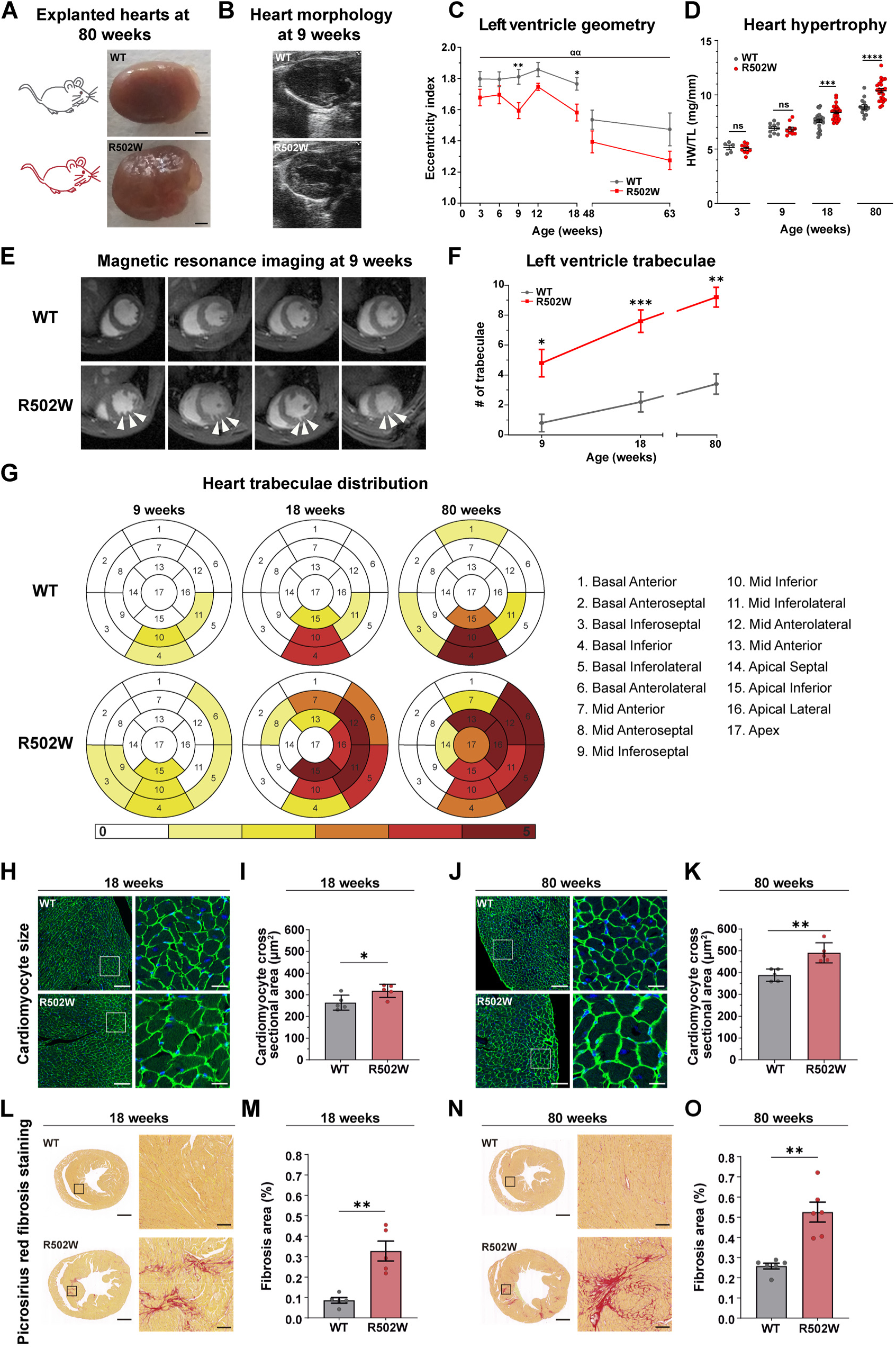
Anatomical and histological alterations of homozygous R502W myocardium. **(A)** Representative explanted hearts from WT and R502W mice. **(B)** Long axis echocardiography view of WT and R502W hearts. **(C)** Left ventricle eccentricity index measured by echocardiography in WT and R502W mice (n = 5 – 10 per genotype and age). **(D)** Atria-free heart weight (HW) to tibia length ratio (TL) ratio in WT and R502W mice (n = 5 – 26 per genotype and age). **(E)** Representative magnetic resonance images from WT and R502W hearts. Arrowheads point to trabeculae in the left ventricle of mutant mice. **(F)** Longitudinal study of the number of trabeculae in the left ventricle of WT and R502W mice (n = 5 animals per genotype). **(G)** Bull’s eye plots of the degree of trabeculation in different ventricular areas. **(H)** Representative images of laminin immunohistochemical staining from sections of 18-week-old WT and R502W hearts used to quantify cardiomyocyte cross sectional area (CSA). **(I)** CSA from 18-week-old WT and R502W animals (n = 5 animals per genotype). **(J)** Representative images of laminin immunohistochemical staining from sections of 80-week-old WT and R502W hearts. **(K)** CSA from 80-week-old WT and R502W animals (n = 5 animals per genotype). **(L)** Representative images of transversal sections of 18-week-old WT and R502W hearts stained with Picrosirius red to detect collagen. **(M)** Percentage of fibrosis area detected in 18-week-old WT and R502W hearts (n = 5 animals per genotype). **(N)** Representative images of transversal sections from 80-week-old R502W and WT hearts stained with Picrosirius red. **(O)** Percentage of fibrosis area detected in 80-week-old WT and R502W hearts (n = 5 animals per genotype). Scale bars are 1 mm in panels A, L and N (100 µm in insets), and 100 µm in panels H and J (20 µm in insets). Insets are indicated by white (H, J) or black (L, N) boxes in images on the left of the panels. Histological sections in this figure were embedded in paraffin. *p < 0.05, **p < 0.01, ***p < 0.001, ****p < 0.0001, ^αα^p < 0.01.

To assess if the tissue alterations in R502W mice affect the pumping activity of the left ventricle, we monitored cardiac function by echocardiography. Mutant mice develop diastolic dysfunction as detected both by analysis of E/A ratios (**Supplementary Figure S3A,B**) and by a ∼50% increase in isovolumic relaxation times (IVRT), which is already evident at 3 weeks of age (**Figure 3A**). At later time points, we also observed a ∼20% reduction in ejection fraction indicative of impaired systole in mutant mice (**Figure 3B**, source ventricular dimensions in **Supplementary Figure S3C,D**). We further confirmed systolic dysfunction in R502W hearts by MRI (**Supplementary Figure S4**).

**Figure 3.**
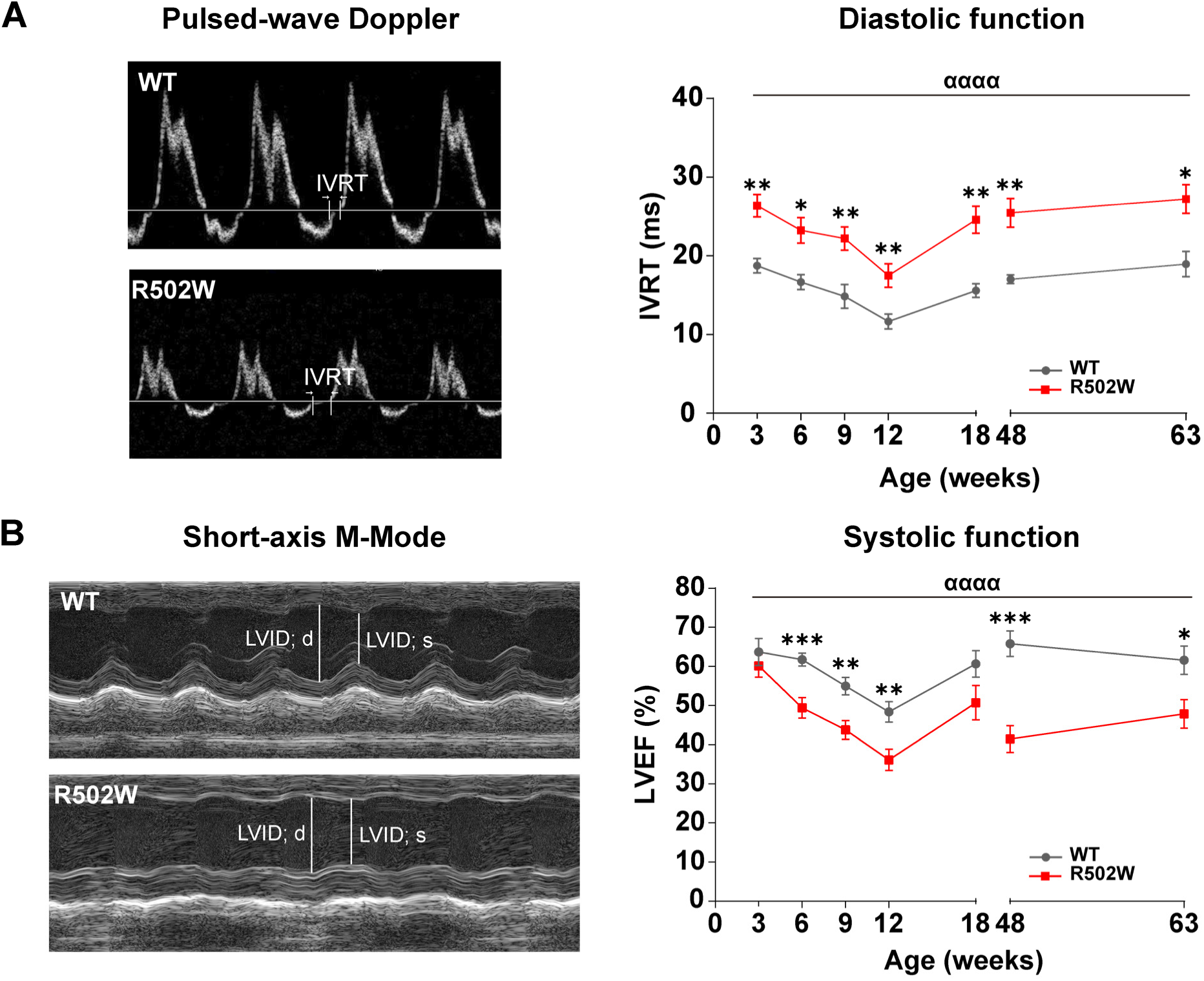
Cardiac dysfunction in R502W mice. **(A)** Isovolumic relaxation times (IVRT) in R502W and WT mice (n = 4 – 19 per each genotype and age). Representative pulsed-wave Doppler measurements of 18-week-old WT and R502W hearts are shown. **(B)** Left ventricle ejection fraction (LVEF) in R502W and WT animals (n = 5 – 20 per each genotype and age). Representative short-axis M-mode acquisitions of 18-week-old WT and R502W hearts are shown. *p < 0.05, **p < 0.01, ***p < 0.001, ^αααα^p < 0.0001.

In summary, anatomical, histological and functional analyses demonstrate that homozygous R502W animals progressively develop several hallmarks of HCM remodelling, indicating that pathogenic mechanisms of variant *MYBPC3* p.R502W are conserved in human and mice. Similar to other mouse models carrying mutant *Mybpc3* genes, cardiac alterations in heterozygous R502W animals are less recognisable if present at all (**Supplementary Figure S5**) ^27^.

### Evolving HCM transcriptional landscape in R502W hearts

Upregulation of stress and fibrosis-related genes as well as altered expression of sarcomeric protein isoforms and metabolic enzymes have been detected both in human myocardium in advanced HCM states ^28–31^ and in murine models of the disease ^18,32^. Considering the progressive HCM phenotype of R502W animals, we exploited RNAseq to examine how gene expression evolves between 3-week-old animals, whose hearts are not hypertrophied and have not developed systolic dysfunction yet, and 18-week-old counterparts, which show an overt HCM phenotype (**Supplementary File S1)**. Principal component analysis indicates that the transcriptional landscapes of the four experimental groups are well separated (**Figure 4A**). There are 970 differentially expressed genes (DEG) between 3-week-old WT and R502W animals, while the number of DEG increases to 1345 in 18-week-old animals (**Figure 4B**). Enrichment pathway analysis showed that DEG with |log_2_ fold change(FC)|≥1 mostly fall in terms related to hypertrophy of the heart and fibrosis, especially in 18-week-old hearts (**Figure 4C-F**). We obtained similar conclusions by analysing the myocardial proteome of 13-week-old mice, in agreement with observations in human HCM ^29^ (**Supplementary Figure S6, Supplementary File S2**). Remarkably, R502W myocardium shows altered transcription of 27% of genes in a list of 329 HCM genes (**Figure 4G,H**) and 52% of the most differentially expressed genes in HCM mouse models caused by mutations in myosin ^18^ (**Supplementary Figure S7**). Both observations indicate that R502W animals develop typical HCM transcriptome. Indeed, mutant hearts show increased transcription levels of stress markers *Nppa* and *Nppb* and genes involved in extracellular matrix remodeling (*Tgfb3*, *Pdgfc, Col3a1, Mmp2, Postn*) (**Supplementary Figure S8A,B**). R502W hearts also express sarcomeric protein isoforms typical of the fetal gene program that is common in HCM (reduced *Myh6*/*Myh7* ratio and increased *Acta1*) ^28,33^ (**Supplementary Figure S8C**). We also detected alteration of expression other sarcomere components, including upregulation of the Z-disc-associated *Csrp3*, *Flnc* and *Des*, which play roles in mechanosensing and structural integration with the non-sarcomeric cytoskeleton ^34^ (**Supplementary Figure S8D**). Regarding metabolic genes, the expression profile of R502W myocardium includes downregulation of *Cpt2*, *Acadvl*, *Hadha, Hadhb, Eci2* and *Ech1*, involved in fatty acid oxidation, and *Pcx*, *Cs* and *Sucla2*, which participate in the tricarboxylic acid cycle (**Supplementary Figure S8E,F**). We also detected downregulation of *Ndufs6*, *Etfdh*, *Uqcrc1*, *Ndufv2* and *Sdha*, which encode proteins of the mitochondrial electron transport system, and several antioxidant enzymes (*Sod1, Cat and Xdh*) (**Supplementary Figure S9A,B**). Different from results with human myocardium, R502W hearts show only minor changes in the expression of genes involved in glucose metabolism ^30^ (**Supplementary Figure S9C**). R502W myocardium also has increased expression of *Ankrd1*, *Fhl1*, *Creb5*, *Xirp2* and *Bdnf*, which have also involved in HCM progression ^31,32,35,36^, and *Kif5b* and *Trim54*, which agrees with the emerging role of the microtubule cytoskeleton in HCM pathophysiology ^37^ (**Supplementary Figure S9D**). Consistent with results obtained with cMyBP-C-null mice, the expression of several genes codifying for potassium channels is altered in R502W animals ^32,38^ (**Supplementary Figure S9E**). Interestingly, we also detected dysregulation of *Pln*, *Ryr2*, *Jph2* and *Cacna1c*, key genes involved in modulation of cytosolic calcium levels, as well as downregulation of *Fhod3*, a recently described cMyBP-C interactor ^39^ (**Supplementary Figure S9F,G**).

**Figure 4.**
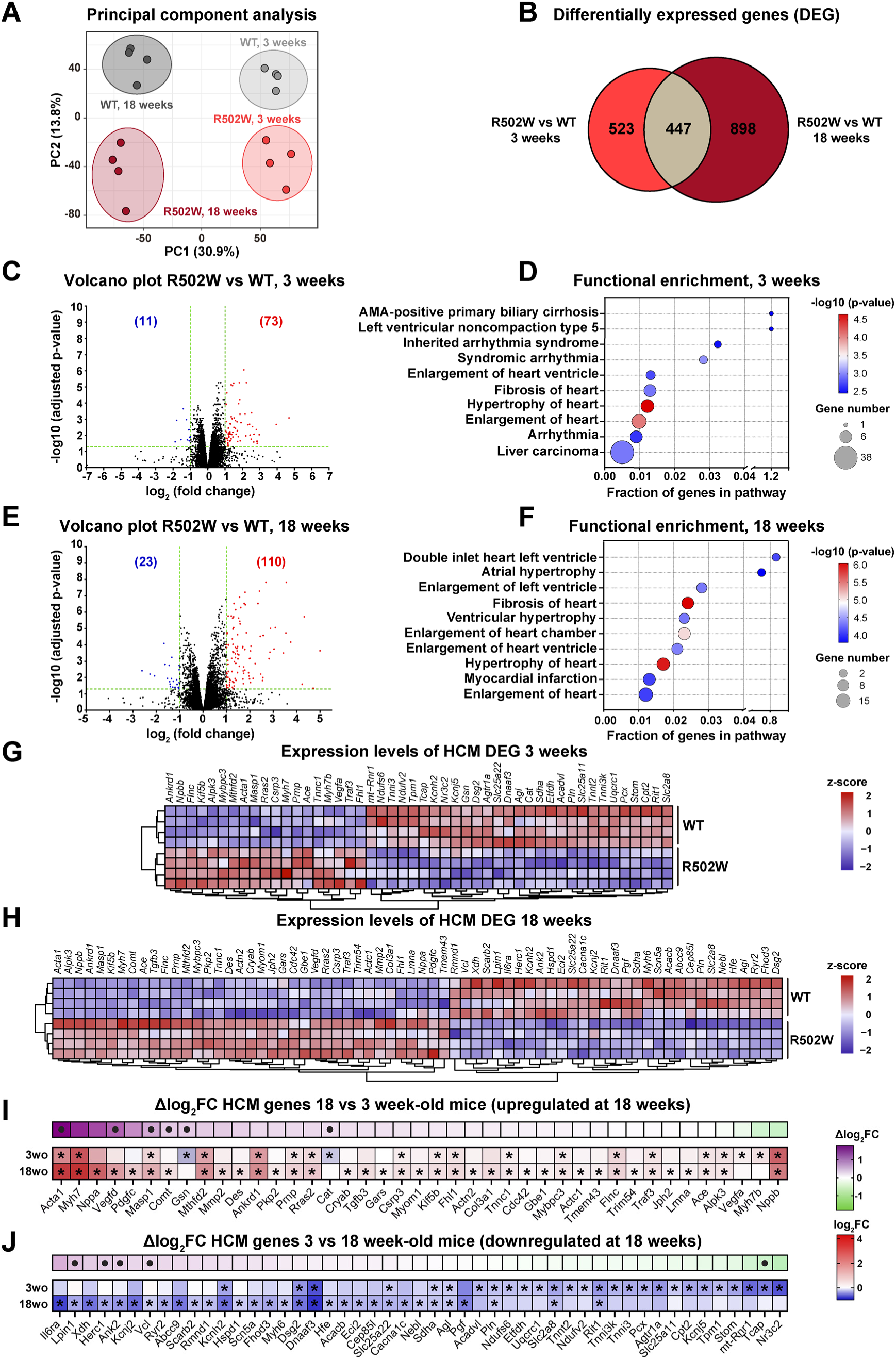
HCM transcriptomic landscape in R502W hearts. **(A)** Principal component analysis from RNAseq data (n=4 per each genotype and age). **(B)** Venn diagram showing the number of differentially expressed genes (DEG) in R502W *vs* WT mice in 3- and 18-week-old animals. **(C)** Volcano plot of gene expression in 3-week-old R502W mice compared to WT. Significantly up- or downregulated genes with |log_2_FC| ≥ 1 are shown in red and blue, respectively. **(D)** Bubble plot representing functional enrichment analysis in toxicology terms for DEG with |log_2_FC| ≥ 1 in 3-week-old animals. **(E)** Volcano plot of gene expression in 18-week-old R502W mice compared to WT. Significantly up or downregulated genes with |log_2_FC| ≥ 1 are shown in red and blue, respectively. **(F)** Bubble plot representing functional enrichment analysis in toxicology terms for DEG with |log_2_FC| ≥ 1 in 18-week-old animals. **(G)** Cluster analysis of HCM DEG in 3-week-old R502W and WT animals. **(H)** Cluster analysis of HCM DEG in 18-week-old R502W and WT animals **(I,J)** Comparative heatmap representing DEG in 3- or 18-week-old animals, and the log_2_FC difference (Δlog_2_FC) between both time points. Genes in panels I and J are upregulated or downregulated, respectively, in 18-week-old animals. In the case of panels I and J, * mark DEG with adjusted pvalue < 0.05 in 3- or 18-week old animals and • represent genes whose Δlog_2_FC have associated adjusted pvalue < 0.05.

Our RNA-seq data indicate that the HCM transcriptional landscape in R502W myocardium is qualitatively similar in 3-and 18-week-old animals, although it intensifies with age (**Figure 4G-J**). Indeed, exacerbated upregulation is apparent for *Acta1*, whose expression is doubled at 18 weeks of age, and *Vegfd*, a key regulator of angiogenesis in the myocardium (**Figure 4I, Supplementary Figures S8C, S10A**). Conversely, downregulation of *Lpin1*, a global modulator of lipid metabolism, and *Ank2* and *Vcl*, encoding scaffolding proteins fundamental for electrical conduction in the myocardium, is only apparent at 18 weeks of age (**Figure 4J, Supplementary Figure S10B**). Intriguingly, *Gsn*, encoding the actin-depolymerizing protein gelsolin, *Cat* and *Tcap*, which codes for the structural component of the sarcomere telethonin, were only transiently downregulated at 3 weeks (**Figure 4I, J, Supplementary Figures S9B, S10C**).

In summary, changes in gene expression of R502W hearts are consistent with data from other mouse models of HCM and from samples of human patients. Remarkably, the HCM transcriptional state of R502W mice is already present at 3 weeks, well before the hypertrophic phenotype is fully overt.

### Preserved expression and localization of cMyBP-C R502W

We tested our initial hypothesis that HCM remodeling in R502W mice occurs in the absence of cMyBP-C haploinsufficiency. To this aim, we measured the expression of *Mybpc3* at the mRNA and protein levels in WT and mutant mice. Despite indication of slight upregulation of *Mybpc3* R502W mRNA in RNAseq (**Supplementary Figure S8D**), quantitative PCR (qPCR) analysis shows no evidence of differential expression of *Mybpc3* transcripts in 3-and 18-week-old animals (**Figure 5A**). Importantly, the levels of cMyBP-C protein are also preserved in 9- and 13-week-old R502W hearts, as assessed respectively by western blot (**Figure 5B**) and proteomic analyses (**Supplementary Figure S11A**). Furthermore, using immunocytochemistry on neonatal cardiomyocytes we observed that both WT and R502W cMyBP-C proteins appear as a regular pattern of doublets, in agreement with the presence of two C-zones per sarcomere (**Figures 1B**, **5C**). No noticeable difference between WT and R502W localization could be detected in fluorescence intensity profiles of cMyBP-C, strongly suggesting that the mutation does not affect incorporation of cMyBP-C into the sarcomere (**Figure 5D,E**). Since the modulatory activity of cMyBP-C depends on phosphorylation ^7^, we also studied whether R502W results in altered phospho-cMyBP-C levels. However, we could not find any differences in the phosphorylation of cMyBP-C R502W (**Supplementary Figure S11B-D**). These experiments also showed that phosphorylation levels of other sarcomere proteins including troponin T and troponin I are preserved in mutant mice (**Supplementary Figure S11E-F**). Overall, our results show that the HCM phenotype in R502W mice develops despite no noticeable alteration of the levels, localization and phosphorylation state of cMyBP-C.

**Figure 5.**
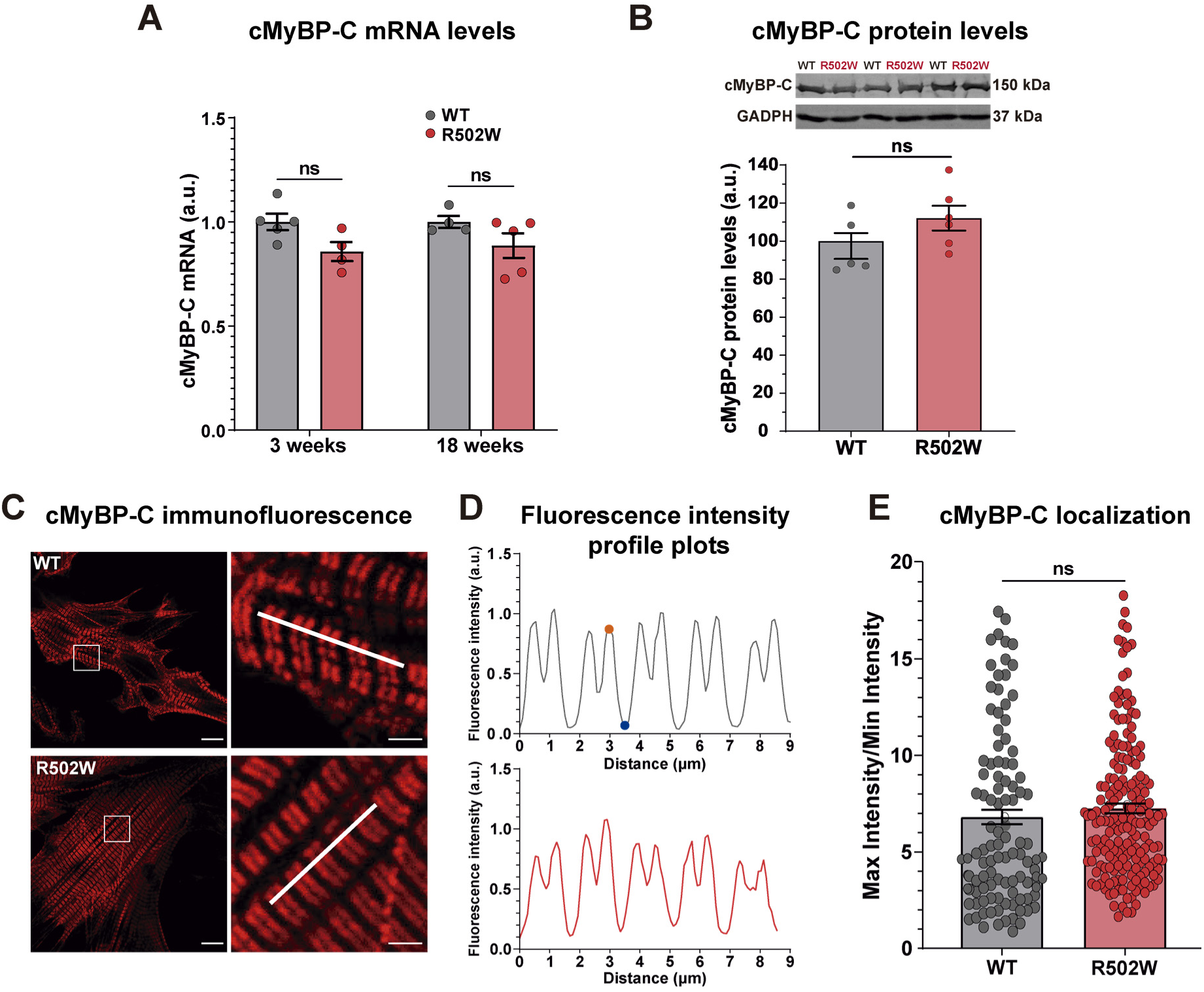
Preserved cMyBP-C levels and localization in R502W mice. **(A)** cMyBP-C mRNA expression levels in the myocardium at 3 and 18 weeks in R502W and WT mice, as measured by qPCR (n= 4 - 5 animals per genotype and age). **(B)** GADPH-normalized cMyBP-C protein levels in the myocardium from 9-week-old mutant and WT mice (n= 5 animals per genotype). **(C)** Sarcomere localization of cMyBP-C in WT and R502W neonatal cardiomyocytes by immunofluorescence. Scale bars are 10 µm (left) add 2 µm (right, zoom region indicated in the left panels by dotted boxes). **(D)** Fluorescence intensity profile plots corresponding to the regions indicated in panel C by white lines. Dots indicate position of maximum (orange) and minimum (cyan) intensity peaks, corresponding respectively to C-band and Z-line. **(E)** Ratio of maximum to minimum intensity peaks from fluorescence intensity plot profiles like the ones presented in panel D (n= 107 ratios from n= 6 cardiomyocytes isolated from 7 WT pups and n= 183 ratios from n= 8 cardiomyocytes isolated from 4 R502W pups). ns: non-significant.

### cMyBP-C R502W binds myosin with less affinity but does not alter SRX/DRX ratio

Considering that cMyBP-C R502W properly incorporates into the sarcomere to the same levels as WT, we examined whether the mutation could alter the modulatory function of cMyBP-C. Since R502W targets the C3 domain of the protein, which has been recently shown to interact with myosin ^24,40^, we tested if R502W could affect this interaction resulting in perturbed myosin SRX/DRX conformations. We first used microscale thermophoresis (MST) to measure binding affinities between recombinant constructs of human β-cardiac myosin and cMyBP-C domains including or not mutation R502W ^41^. MST results indicate that R502W induces 30-50% reduced binding affinity between myosin and c-MyBP-C (**Figure 6A**). Next, we measured myosin SRX/DRX ratios in permeabilized left ventricle tissue from 18-week-old WT and R502W mice using the rate of exchange of fluorescent mant-ATP in the active site of myosin (**Figure 6B,C**) ^9,14^. We also included samples from homozygous cMyBP-C KO mice ^42^, which show the expected faster exchange of mant-ATP as compared to WT indicating a lower proportion of myosin molecules in the SRX state (**Figure 6D,E**). In contrast, the decay curves for R502W and WT overlap to a great extent (**Figure 6D**). Indeed, the estimated proportion of myosin molecules in the SRX state is very similar for WT and R502W tissues (**Figure 6E**). Hence, the observed reduced interaction between myosin and cMyBP-C R502W does not translate into noticeable alteration of the SRX/DRX conformations of myosin in R502W myocardium.

**Figure 6.**
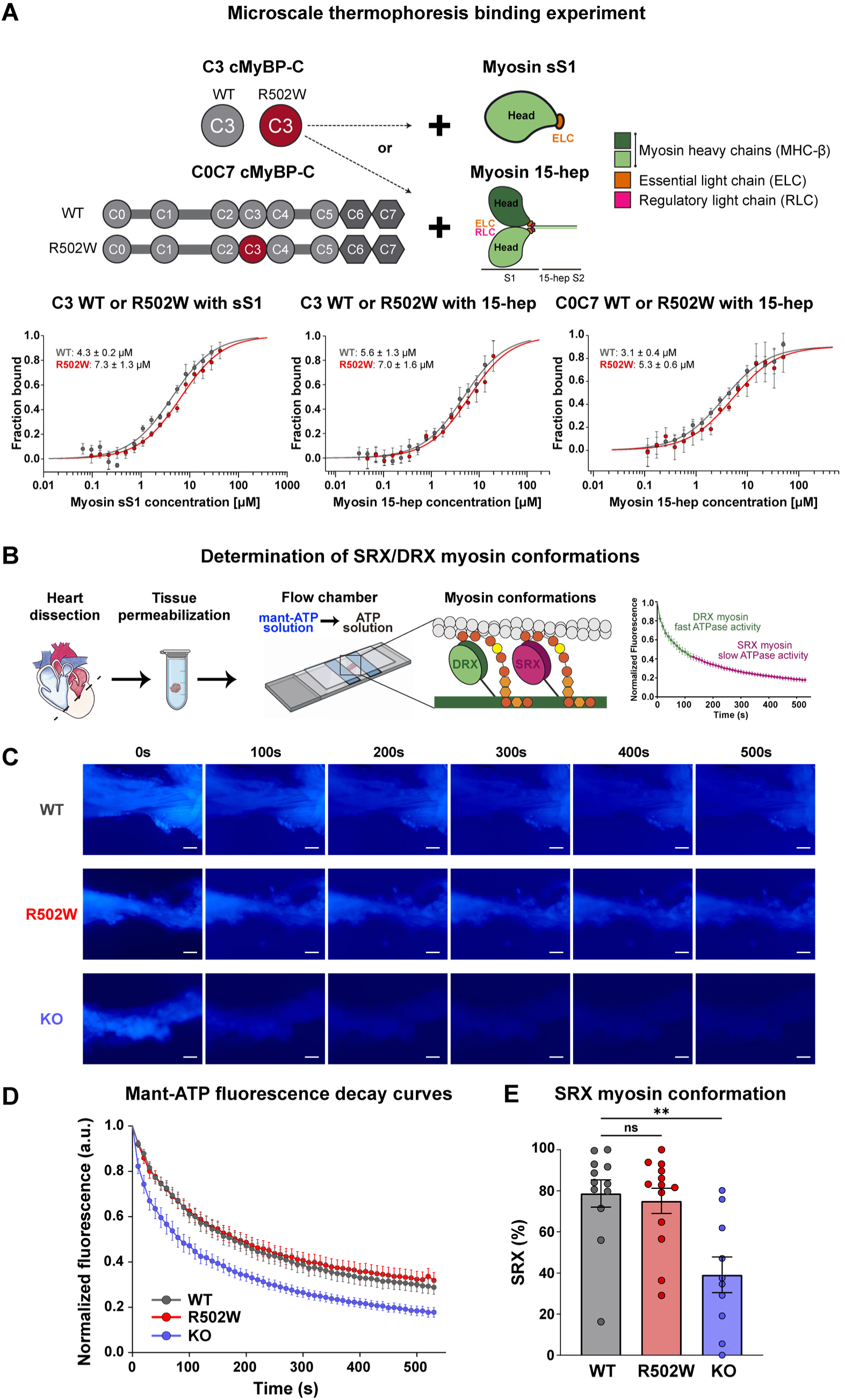
R502W decreases the affinity of cMyBP-C for myosin but does not alter the proportion myosin’s active conformations. **(A)** Effect of R502W mutation on the interaction between cMyBP-C C0C7 or C3 domains with recombinant cardiac myosin sS1 or 15-hep as assessed by MST. Graphs show the average of n = 5 – 8 replicate runs from a single set of protein preparations per plot. Solid lines represent binding isotherms using a monovalent binding model. Mean dissociation constants are indicated. **(B)** Diagram of the fluorescence-based mant-ATP experiment to quantify SRX and DRX states of myosin in permeabilized cardiac tissue. **(C)** Representative images showing fluorescence intensity decay in mant-ATP experiments using samples from 18-week-old WT, R502W and KO mice. Scale bars are 100 µm. **(D)** Average man-ATP fluorescence decay curves from WT, R502W and KO myocardium (n = 10 – 13 experiments using n= 5 – 6 animals per genotype). **(E)** Percentage of SRX myosin in WT, R502W and KO myocardium. **p < 0.01; ns: non-significant. The figure includes elements from BioRender.com.

### Mavacamten is beneficial for cMyBP-C R502W and KO mice

Mavacamten has been shown to normalize SRX/DRX myosin conformations in cMyBP-C haploinsufficient HCM myocardium, an effect that has been linked to the therapeutic benefit of the drug^16^. Hence, we hypothesized that patients carrying missense variants in *MYPBC3* that do not affect cMyBP-C levels nor cause changes in myosin SRX/DRX ratio may benefit less from mavacamten. To explore this possibility, we treated 3-week-old WT, R502W and cMyBP-C KO mice with the drug for 21 weeks (**Figure 7A**). Four weeks after starting the treatment all treated groups had therapeutic plasma levels of mavacamten (**Supplementary Figure S12A**) ^20^. Remarkably, results show that mavacamten prevents the development of cardiomyocyte hypertrophy (**Figure 7B,C**) and myocardial collagen deposition (**Figure 7D,E**) both in R502W and KO hearts. We also found a tendency to less heart weight in mavacamten-treated R502W mice despite the low number of animals tested (**Supplementary Figure S12B**). In agreement with previous results with other HCM mouse models ^18^, mavacamten appears to have limited ability to revert alterations present before treatment. Specifically, the drug does not normalize left ventricle eccentricity in neither HCM model (**Supplementary Figure S12C-E**) or number of ventricular trabeculae in R502W animals (**Supplementary Figure S12F-H**). Regarding functional cardiac parameters, we found that mavacamten induces a modest decrease in IVRT in treated R502W animals; however, the same tendency is also observed in WT mice, suggesting that this effect on diastolic function could derive from myosin inhibition directly (**Supplementary Figure S12I-K**). Similarly, as expected, we observed that mavacamten decreases the left ventricular ejection fraction of wild-type animals ^18^. This effect was not so evident in the HCM groups, probably reflecting convolution between the mechanism of action of the drug and its ability to block HCM progression (**Supplementary Figure S12L-N**). To examine how these seemingly opposed effects emerge at the organismal level, we subjected the six experimental groups to effort tests (**Figure 7F**). We found that mavacamten decreases the exercise capacity of WT animals (**Figure 7G**), in agreement with the observed reduction in ejection fraction. Interestingly, treatment with mavacamten resulted in improved exercise capacity of R502W mice, indicating an overall beneficial effect of the drug in these animals (**Figure 7G**). In contrast, the exercise capacity of KO mice treated with the drug did not improve (**Figure 7G**). In summary, our results show that mavacamten blunts progression of key aspects of HCM remodeling in R502W and KO mice, leading to better exercise capacity in R502W animals.

**Figure 7.**
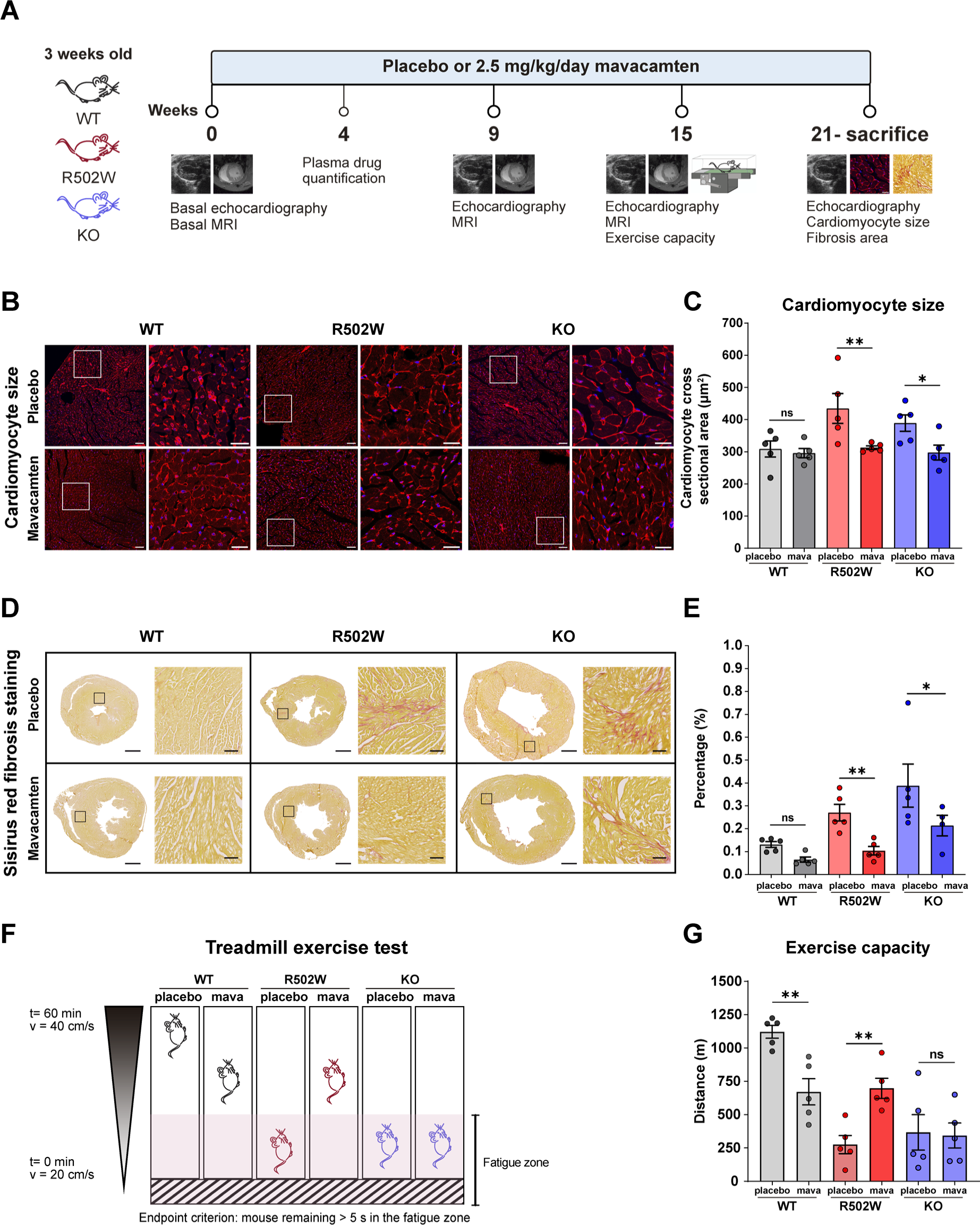
Treatment with mavacamten blunts hypertrophic remodeling in R502W and KO hearts. **(A)** Diagram of the experimental design to test mavacamten efficacy. **(B)** WGA immunostaining to quantify cardiomyocyte cross sectional area from WT, R502W and KO tissue treated or not with mavacamten. **(C)** Cardiomyocyte cross sectional area in WT, R502W and KO animals treated or not with mavacamten (n = 5 animals per genotype). **(D)** Picrosirius red staining of myocardial sections from WT, R502W and KO mice treated or not with mavacamten. **(E)** Myocardial fibrosis area from WT, KO and R502W mice treated or not with mavacamten (n = 4-5 animals per genotype). **(F)** Schematic representation of the treadmill-based exercise capacity test. Speed was increased from 20 cm/s to 40 cm/s in 60 minutes according to the protocol shown on the left. **(G)** Maximum distance run before reaching the endpoint criterion for the three genotypes treated or not with mavacamten for 15 weeks (n = 5 animals per genotype). Scale bars are 100 µm in panel B (50 µm in insets), and 1 mm in panel D (100 µm in insets). Insets are indicated by white (B) or black (D) boxes in images on the left of the panels. Histological sections in this figure were embedded in OCT. *p < 0.05, **p < 0.01, ns: non-significant.

## DISCUSSION

Clinical management of HCM has improved considerably over the last three decades thanks to the identification of causative genes leading to better diagnosis and pointing to pathomechanisms that are now the target of innovative therapies ^2^. For instance, building on the observation that the disease typically debuts as a hypercontractile myocardium, and that many patients carry gain-of-function variants in *MYH7*, myosin inhibitors have been developed as candidate therapies ^19^. In the EXPLORER-HCM clinical trial, mavacamten, the first-in-class of this family of drugs, resulted in improved cardiac function, exercise capacity and health status in patients with symptomatic obstructive HCM ^20^. These positive results prompted the FDA to approve mavacamten for the treatment of this condition in April 2022, a paradigm shift in the management of a disease that had been orphan of specific treatments ^5^.

Notwithstanding these advances, the heterogenous genetics of HCM patients still pose major challenges that remain unmet ^8,43,44^. In particular, it is possible that the response to therapeutic approaches varies across different HCM-causing variants, which may contribute to the limited efficacy of mavacamten in a fraction of patients ^20,45^. In this regard, several lines of evidence indicate that both variants in *MYH7* and in *MYBPC3* (provided they induce cMyBP-C-haploinsufficiency) decrease myosin SRX/DRX ratios in the myocardium ^14–16,22^, a phenotype that is normalized by mavacamten *in vitro* ^16,22^. These results suggest that the drug can have similar therapeutic benefit in HCM caused by *MYH7* variants or by cMyBP-C haploinsufficiency, a scenario supported by the efficacy of mavacamten in HCM mouse models carrying mutations in myosin ^18^ and by our results with cMyBP-C KO mice (**Figure 7**).

Much less is known about primary pathomechanisms triggered by *MYBPC3* variants that do not cause protein haploinsufficiency, which is problematic because they are responsible for a substantial fraction of HCM cases ^7,23,26^. To the best of our knowledge, the R502W strain is the first murine model for this type of variants. Our result that the R502W mutation does not lead to cMyBP-C haploinsufficiency, which has also been observed in cardiomyocytes derived from human induced pluripotent stem cells ^46^, agrees with the subtle effects of the arginine-to-tryptophan substitution in the structure and stability of the parent C3 domain of cMyBP-C ^23,26,47^. In addition, we have verified that the mutant protein is properly incorporated into cardiac sarcomeres of R502W mice. Despite these normal cMyBP-C levels and localization, R502W animals develop progressive HCM remodeling including cardiac and cardiomyocyte hypertrophy, typical HCM cardiac transcriptome, myocardial interstitial fibrosis, hypertrabeculation, and altered left ventricle geometry and systolic and diastolic dysfunction (**Figures 2-5**). Prompted by these results, we tested if mavacamten is equally effective to halt HCM progression in R502W and cMyBP-C KO mice despite their different underlying primary pathomechanisms, and found that the drug blunts cardiomyocyte hypertrophy and myocardial fibrosis similarly in both cases (**Figure 7**). Hence, our data and evidence in the literature strongly suggest that mavacamten is equally effective for carriers of any pathogenic variant in *MYH7* or *MYBPC3*. Intriguingly, despite the fact that both R502W and KO animals respond to mavacamten at the histological level, only R502W mice improve tolerance to exercise (**Figure 7G**). We interpret that this differential effect stems from the more aggressive phenotype of KO mice (**Supplementary Figure S12B**), which is obvious already in 3-week-old animals ^48^. These results suggest that prompt treatment could boost mavacamten’s efficacy by preventing HCM remodeling, whereas full reversal of established pathogenic phenotypes may be more challenging (**Supplementary Figure S12 C-H**) ^18^. However, the detrimental effect of mavacamten on the exercise capacity of healthy WT animals (**Figure 7G**) reinforces the need for careful evaluation of the risk/benefit balance of preventive treatment with mavacamten ^5^.

Remarkably, while the effects of mavacamten in cMyBP-C KO animals are most probably contributed by normalization of SRX levels ^16^, the benefits of the drug in R502W mice occur in the absence of cMyBP-C haploinsufficiency or basal alteration of SRX/DRX myosin conformations (**Figures 5**, **6D**), indicating that dysregulation of myosin active conformations is not a universal pathomechanism in *MYBPC3*-associated HCM. The observation that R502W reduces the binding affinity between myosin and cMyBP-C (**Figure 6A**) suggests that the pathogenic nature of mutant cMyBP-C may involve alteration of other mechanical properties of myosin ^49^ and that mavacamten may be able to counter these deleterious effects either directly or indirectly, for instance by increasing SRX/DRX ratios above physiological levels as observed in WT myocardium ^16^. Our mechanistic results also have implications for alternative therapeutic strategies in HCM. For example, gene therapies based on expression of cMyBP-C ^50^ may show less efficacy in carriers of variants like R502W if only limited dilution of the poison-peptide effect of the mutant is achieved. Similarly, the benefits of exon skipping approaches should be carefully evaluated if they result in complete loss of important cMyBP-C-myosin interactions ^51^. Undoubtedly, base editing emerges as a highly attractive therapeutic approach in HCM caused by single-base substitutions in cMyBP-C as in the case of variants in *MYH7* ^52,53^.

The modulatory role of cMyBP-C on sarcomere contraction is complex and not fully understood ^7^. The C-terminal domains of the protein are strongly bound to the thick filament through strong electrostatic interactions with different regions of myosin molecules ^54,55^ whereas the N-terminal domains are more dynamic and capable of regulating contraction by reversible interactions that stabilize the active state of thin filaments and the inactive state of thick filaments ^56^. The central C3-C7 domains of cMyBP-C have traditionally been regarded as mere connectors of the N- and C-terminal regions of the protein (**Figure 1B**); however, this apparently minor functional role of C3-C7 domains does not agree with the fact that many missense HCM variants, including R502W, target these domains ^23,26,57^. Indeed, recent reports have found that the central region of cMyBP-C interacts with myosin, as assessed both biochemically ^24,40^ and structurally ^54^. Although further work is needed to unravel specific molecular mechanisms of myosin modulation by central domains of cMyBP-C, our findings support the functional relevance of interactions of myosin and C3-C7 cMyBP-C domains, as well as the proposal that alteration of myosin-cMyBP-C interactions can be a molecular pathomechanism in HCM ^40^. Enticingly, the vast majority of candidate pathogenic, non-protein-haploinsufficient variants targeting central domains of cMyBP-C (R495W, R495G, R495Q, R502Q, R810H, R820W, R820Q) ^23^ affect the surface charge of targeted domains in a similar manner to R502W. This is a typical mechanism of disruption of protein-protein interactions ^47^, which suggests that alterations of surface electrostatics could be used to pinpoint rare pathogenic variants in *MYH7* and *MYBPC3* for which strong clinical cosegregation is not available and that would be otherwise classified as of uncertain significance ^23^.

Unlike the severe cardiac hypertrophy of fully cMyBP-C haploinsufficient mouse models, the relatively mild cardiac phenotype of R502W mice is more similar to how HCM typically manifests in human hearts^27,58^. Strikingly, most key features of HCM cardiac transcriptome identified in advanced human disease and other HCM mouse models are already present in 3-week-old R502W mice, suggesting that sarcomere defects induced by pathogenic variants rapidly induce lasting transcriptomic alterations that precede full establishment of cardiac dysfunction and hypertrophy. Similar results have been obtained in postnatal cMyBP-C null mice ^32^. We identified one notable exception to this general trend; while it is well established that patients with advanced disease show global alterations of cardiac energetics and metabolism including downregulation of both lipid and glucose metabolism ^30^, R502W transcriptional changes are more restricted to lipid pathways. This observation suggests that changes in glucose metabolism in advanced HCM are secondary to other alterations during disease progression.

In summary, our preclinical data indicate that all carriers of pathogenic variants in *MYBPC3*, arguably the most common cause of HCM ^7^, may benefit from myosin inhibition similarly regardless of diverging specific primordial pathomechanisms (**Figure 8**). Our results do not exclude the possibility that variants in other HCM genes may respond differently. However, considering that myosin inhibition appears equally beneficial for carriers of *MYH7* or *MYBPC3* variants, which in combination account for ∼80% of genotype-positive HCM individuals ^7^, the reported limited efficacy of mavacamten in a large fraction of patients is probably due to reasons other than the specific HCM variants they carry, possibly including environmental contributions or other genetic factors unrelated to primary sarcomere deficits. Mechanistically, our results identify altered cMyBP-C-myosin interactions at the basis of pathogenicity of *MYBPC3* variants, an information that can be exploited to improve the yield of genetic testing in HCM and underscores the need to better understand how cMyBP-C and myosin cooperation enables healthy contractions of the heart.

**Figure 8.**
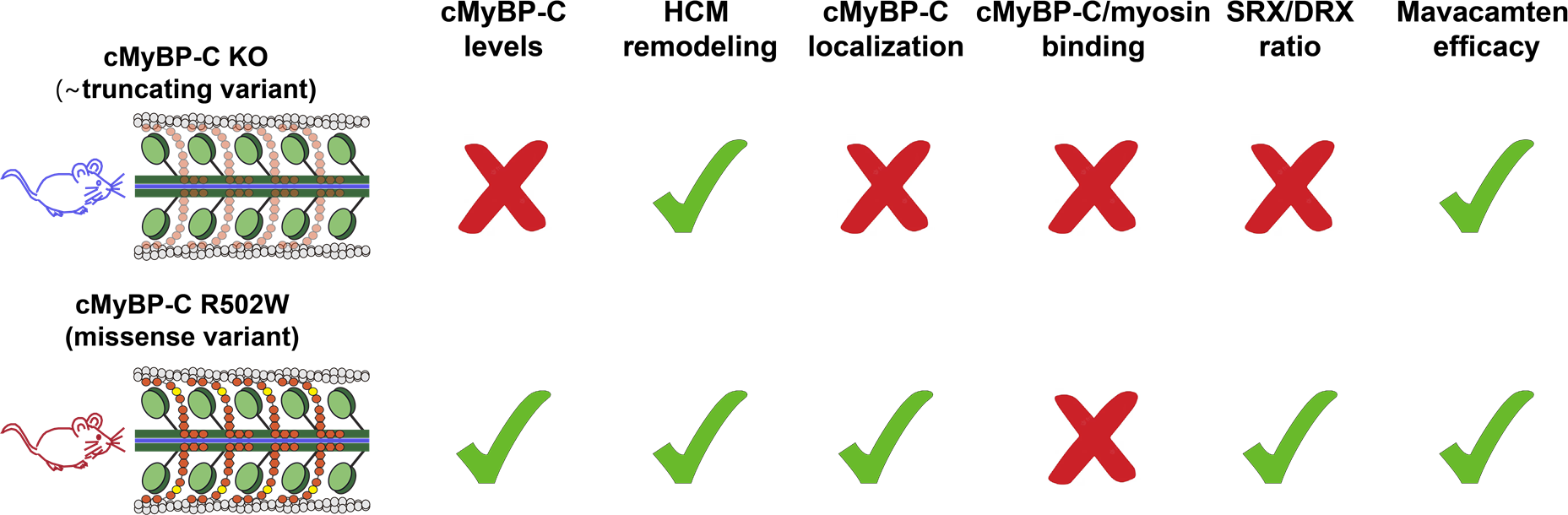
Summary of the main results of this study. We have used cMyBP-C KO mice ^42^ as a model of truncating variants resulting in protein haploinsufficiency, and developed cMyBP-C R502W as a model of missense variants not causing reduced cMyBP-C levels. Despite non-overlapping molecular pathomechanisms, both mice exhibit HCM remodeling. Unlike KO mice and despite altered cMyBP-C-myosin interaction, R502W animals show preserved cMyBP-C localization and SRX/DRX states of myosin. However, both KO and R502W animals benefit from treatment with myosin-inhibitor mavacamten.

## METHODS

### Animal experimentation

All procedures were carried out in accordance with Spanish and European legislation (RD53/2013, Law 32/2007 and Directive 2010/63/EU) and approved by the Madrid Regional Government’s Animal Protection Area (PROEX 042/18 and 107.8/23). Mice were housed in ventilated cages with 12 day/night cycles and access to food and water *ad libitum*.

### Mouse models

CRISPR-Cas9 mediated homology-directed genome editing was used to generate knock-in cMyBP-C R502W animals (**Supplementary Figure S1B,C**). Briefly, superovulated C57BL/6J female mice were crossed with male counterparts and resulting zygotes were collected. Using a micromanipulator, Cas9 protein, designed single-guide RNA (sgRNA) and template single-stranded DNA (ssDNA) were microinjected into the pronucleus of zygotes (**Supplementary Table S1**). Injected zygotes were cultured and two-cell embryos were transferred to pseudo-pregnant C57BL/6J females. Finally, these females were bred to obtain founder (F0) offspring including non-edited and edited mice. Germline transmission of the R502W variant was verified crossing F0 mice with WT C57BL/6J animals. F0 and F1 generations were genotyped using outward and inward primers and by Sanger sequencing (**Supplementary Table S1**). For management of R502W colony, mice were genotyped using rhAmp SNP Genotyping from IDT (**Supplementary Figure S13**, **Supplementary Table S1**). KO mice ^42^ were bred into C57BL/6J background and genotyped with primers in **Supplementary Table S1**.

### Echocardiography

Non-invasive transthoracic echocardiography was performed on anesthetised mice using 1%-2% isoflurane in 100% oxygen, by personnel blinded to the study, using a high-resolution ultrasound imaging system with 30-MHz transducer (Vevo 2100, MS400, VisualSonics,Canada). Standard parasternal long or short-axis views in two-dimensional (2D) mode and M-mode images of the left ventricle (LV) were acquired. Long-axis images were used to estimate LV eccentricity index, which was calculated as the ratio of longitudinal vs transversal LV lengths. M-mode images at the level of the papillary muscles were used to estimate LV end-diastolic and end-systolic volumes (LV Vol;d and LV Vol;s, respectively) and to calculate ejection fraction ((LV Vol;d – LV Vol;s) / LV Vol;d x 100). Diastolic function was evaluated by pulsed-wave Doppler using a 2D apical view to estimate mitral valve inflow.

### Magnetic resonance imaging

MRI studies were performed using a 7-T Agilent/Varian scanner (Agilent, Santa Clara, CA, USA) equipped with a DD2 console. Two-chamber short-axis cardiac MRI was performed using an actively shielded 115/60 gradient. A surface coil was used for signal transmission and reception. MRI experiments were conducted by applying an ECG-triggered fast gradient echo cine sequence with the following imaging parameters: ∼125/1.25 msec TR/TE (minimum values depending on heart rate and number of frames per heart cycle); 30 x 30 mm field of view; 128x128 acquisition matrix; 15^°^ flip angle; 4 averages; 20 cardiac phases; ∼12 slices (exact number of slices are chosen to cover the entire left ventricle); 0.8 mm slice thickness and 0.2 mm gaps. Cine MRI was analyzed using Segment software v1.9 R3819 (http://segment.heiberg.se) ^59^. Epicardial and endocardial borders were delineated manually on the short axis of each slide to measure total ventricular volumes and cardiac mass ^60^. Regarding analysis of trabeculae, for each segment, a value of 1 was assigned if the tissues contained trabeculations; otherwise, the value assigned was 0. The sum of all these values per segment for all animals in the experimental groups was represented in bull’s eye maps.

### Histology

Following sacrifice in a CO_2_ chamber, heart and thoracic arteries were removed and washed with 0.1 M phosphate buffered saline (PBS; pH 7.3). The aorta, pulmonary artery and atria were excised and the cardiac ventricles were fixed with 4% paraformaldehyde (PFA) for 48 h before embedding in paraffin or OCT (Sakura Finetek). For analysis of cardiomyocyte cross sectional area, 8-µm transversal sections from apical to medial myocardial regions were permeabilized with 0.1-0.3% Triton X-100 in PBS for 10 min at room temperature (RT) and blocked with 10% bovine serum albumin (BSA) and 2% normal goat serum (NGS) in PBS during 1 h. Next, sections were labeled with a rabbit anti-mouse laminin primary antibody (Sigma Aldrich, L9393, 1:500 in PBS, 5% BSA, 1% NGS, overnight at 4°C) or, alternatively, wheat germ agglutinin (WGA) conjugated to Alexa Fluor 555 (1:250 in PBS, 5% BSA, 1% NGS, 1.5 h at RT). Preparations were washed three times with PBS for 5 min at RT. In the case of sections stained with laminin, species-matched Alexa Fluor 488 labelled secondary antibody was used (Thermo Fisher W11261, 1:500 in PBS, 5% BSA, 1% NGS, 2 h at RT) followed by three washes with PBS for 5 min at RT. Finally, samples were incubated with 5 µg/mL 4′,6-diamidino-2-phenylindole dihydrochloride (DAPI, Sigma-Aldrich) in PBS, 5% BSA, 1% NGS, 5 min at RT and mounted using Mowiol 4.88 mounting medium. Fluorescence images of Alexa Fluor 488 (laminin immunostaining) and Alexa Fluor 555 (WGA immunostaining) were acquired using a Nikon A1-R confocal microscope with a Plan Apo 40x/1.3 Oil objective. Three images from three different sections per each mouse heart were obtained and analyzed using the open-source deep learning-based segmentation tool Cellpose ^61^ in the Fiji ^62^ distribution of ImageJ with the help of a Cellpose wrapper developed by the BioImaging And Optics Platform (BIOP, Ecole Polytechnique Fédérale de Lausanne, Switzerland). For determination of fibrosis, paraffin or OCT embedded hearts were sectioned transversally at a thickness of 5 µm from apex to base (150 µm separation between sections) to obtain a complete short-axis, two-chamber view. Sections were stained using Picrosirius red staining and images were digitalized using a Hamamatsu digital slide scanner. Quantitative analysis of fibrosis area was performed using Fiji and a custom script written in ImageJ macro language that measures area occupied by red-hued pixels. For each heart, the percent area of fibrosis was obtained from 8 -12 cross-sectional areas.

### Transcriptomics

Cardiac RNA was isolated from 3- and 18-week-old hearts that were stored at −80°C in 500 μL of RNAlater™ Stabilization Solution (Invitrogen, Cat. No. AM7020). Total RNA was isolated by homogenization with T10 basic ULTRA-TURRAX® (IKA), using TRI reagent followed by chloroform precipitation and a purification step using Rneasy micro-Kit (Qiagen, Cat. No. 74004) following the indications of the provider. RNA quality was quantified from its absorbance at 260 nm, while purity was assessed from the ratios of absorbance at 260 nm and 280 nm, and 260 nm and 230 nm. Sample quality was evaluated using RNA Integrity Number (RIN), which was measured using a 6000 Nano chip in a 2100 Bioanalyzer (Agilent Technologies). For next generation sequencing (NGS) experiments, RNA with RIN > 9 were single-end sequenced on a HiSeq4000 platform (Illumina) with a minimum of 8 M reads per sample at a 50 nt read length. NGS experiments were performed in the Genomics Unit of the CNIC. Raw data were pre-processed with a pipeline using cutadapt to remove Illumina adapter and FastQC to perform quality control. RSEM was used to align processed reads against a reference transcriptome derived from GRCm38.99, and to obtain gene-expression levels. The percentage of successfully mapped reads was 87% on average. Differentially expressed genes (DEG, adjusted p-value < 0.05), were estimated using limma analysis of raw counts on a collection of 14,109 genes showing at least 1 count per million in no less than 4 samples. Subsequently, the Benjamini-Hochberg method was used to adjust the p-values and control the false positive rate. We used Ingenuity Pathway Analysis (IPA, QIAGEN) to further analyze RNAseq results. We did not restrict analyses in IPA to specific species, cell type or other characteristics. We used enrichment bubble plots analysis considering DEG with absolute FC ≥ 2 or ≤ 2 (|log_2_FC| ≥ 1) to represent the top 10 enriched IPA toxicological functions. For the representation of HCM heatmaps, we used all DEGs involved in IPA’s HCM pathway. The percentage of DEGs belonging to the HCM pathway present in our model with respect to IPA’s HCM pathway was calculated considering the 329 genes detected in our RNAseq. For differential expression analysis between 3- and 18-week-old animals, we calculated Δlog_2_FC = log_2_FC _18wo_ – log_2_FC _3wo_ for genes upregulated at 18 weeks or Δlog_2_FC = log_2_FC _3wo_ – log_2_FC _18wo_ for those downregulated at 3 weeks. Multiple testing with Benjamini-Hochberg correction was used to determine statistical significance of Δlog_2_FC data. We also calculated the fraction of genes in a list including the most differentially expressed genes in HCM mouse models with Myh6 mutations ^18^ that appeared as DEG in R502W mice. Heatmaps and bubble plots were obtained with software from http://www.bioinformatics.com.cn/srplot.

### qPCR

Reverse transcription was performed starting from 1 μg of template RNA using SuperScript™ IV VILO™ Master Mix (Thermo Fisher Scientific, Cat. No. 11756050) and the following temperature protocol: 10 min at 25 °C, 10 min at 60 °C and 5 min at 85 °C. cDNA samples were diluted 20 times and used for qPCR using Power SYBR™ Green PCR Master Mix (Thermo Fisher Scientific, Cat. No. 4367660) in a Bio-Rad CFX384 thermal cycler using specific primers sets for *Mybpc3* (Forward: TCAAATGGCTGAAGGATGGG, Reverse: GCCTATATGGGACACCTTTATG) and *Actb* as a housekeeping gene (Forward: TGTTACCAACTGGGACGACA and Reverse: GGGGTGTTGAAGGTCTCAAA). The qPCR protocol included denaturation at 95 °C for 10 min, followed by 40 cycles of 15 s at 94 °C and 60 s at 60 °C. The relative gene expression (ΔΔCq) was calculated using the CFX software Maestro™ (BioRad). All RT-qPCRs were performed in technical triplicates.

### Western blot

Hearts were extracted, flash frozen in liquid N_2_, and stored at –80°C. Tissue homogenization was done in an automatic homogeniser (VWR, Pellet Mixer, 47747-370) in 50 mM Tris-HCl pH 6.8, 8M urea, 2M thiourea, 3% sodium dodecyl sulphate (SDS), 10% glycerol, 0.03% SERVA Blue G and 17 mM dithiothreitol (DTT, added freshly). Samples were left in ice for 30 min and then boiled for 3 min while shaking. Finally, they were centrifuged at 18,407 g for 3 min at RT to obtain supernatants. Protein concentration was determined using Bradford assay (Sigma, Cat. No. B6916). 35 μg of protein were electrophoretically separated using 8% SDS-PAGE and transferred to PDVF membranes for 2.5 h at 200 mA per membrane using a semi-dry system at 4°C. Next, membranes were blocked in 5% non-fat milk dissolved in Tris buffer solution (19,5 mM Tris pH 7.6, 150 mM NaCl) containing 0.1% Tween-20 (TBS-T) for 1 h at RT. Membranes were incubated with mouse anti cMyBP-C monoclonal antibody (Santa Cruz Biotechnology, sc-137237, diluted 1:2000 in TBS-T) at 4°C overnight. Next, they were washed for 10 min with TBS-T three times, and incubated for 1h at RT with a chicken anti-mouse antibody conjugated to Alexa Fluor 647 (Life Technologies, Thermo Fisher, diluted 1:5000 in TBS-T). After incubation with the secondary antibody, three more TBS-T washes were performed, and the membrane was incubated for 2h at RT with rabbit anti-GAPDH antibody (Sigma Aldrich, G9545, diluted 1:2000 in TBS-T) at RT. Finally, the excess antibody was removed using 10-min washes with TBS-T and a goat anti-rabbit antibody conjugated with Alexa Fluor 488 (Life Technologies, Thermo Fisher scientific, diluted 1:5000 in TBS-T) was added for 1 h at RT. Results were visualized in an iBright 1500 reader (Invitrogen, Thermo Fisher). cMyBP-C levels normalized to GAPDH were quantified using Quantity One software. Three technical replicates were done.

### Proteomics

Protein cardiac extracts were obtained by tissue homogenization with ceramic beads (MagNa Lyser Green Beads, Roche) in lysis buffer (50 mM Tris-HCl, 2% SDS, pH 6.8) freshly supplemented with 0.1 mM PMSF protease inhibitor. Around 200 μg of protein were subjected to in-filter reduction and alkylation using iodoacetamide followed by trypsin digestion (Nanosep Centrifugal Devices with Omega Membrane-10K, PALL), following the protocol from Wiśniewski et al.^63^. Labelled peptides were injected onto a C-18 reversed phase nano-column (75 μm I.D. and 50 cm, Acclaim PepMap) and separated in a continuous acetonitrile gradient consisting of 8-31% B-solution (0.1% formic acid (v/v) in acetonitrile) for 240 min, and 50-90% B for 1 min, at a flow rate of ∼200 nL/min, using a UPLC-Ultimate 3000 chromatography system connected to a Q-Exactive HF mass spectrometer (Thermo Fisher). Mass spectra were acquired in a data-dependent manner, with an automatic switch between MS and MS/MS using a top 20 method. MS spectra were acquired in a 390-1700 m/z range at 60,000 FT resolution. HCD fragmentation was performed at 33 of normalized collision energy and MS/MS spectra were analysed with a resolution of 30,000. Dynamic exclusion was set to 40 s. For peptide identification, MS/MS spectra were searched using the SEQUEST HT algorithm implemented in Proteome Discoverer 2.5 (Thermo Scientific) against the Uniprot database (mouse_202105_uni-sw.target-decoy.fasta) ^64^. Trypsin digestion was employed with a maximum of 2 missed cleavages. Fixed modifications included Cys carbamidomethylation (57.021464 Da) and TMT labelling at the N-terminal end and Lys (229.162932 Da), with Met oxidation (15.994915 Da) considered as a dynamic modification. Precursor mass tolerance was set at 800 ppm and fragment mass tolerance at 0.03 Da. The precursor charge range was set to 2-4. The false discovery rate (FDR) was calculated using the target/decoy strategy, employing the corrected XCorr score (cXCorr) ^65,66^, along with an additional filter for precursor mass tolerance of 15 ppm ^67^. A 1% FDR was employed as the criterion for peptide identification. Quantitative information was extracted from TMT reporters ions in the MS/MS spectra, and protein abundance changes were analysed using the WSPP model ^68^ by applying the Generic Integration Algorithm ^69^, using the iSanXoT program ^70,71^. In this model, quantitative protein values are expressed using the standardized variable Zq (i.e., normalized log2-ratios expressed in units of standard deviation according to the estimated variances). Median Zq values from R502W and WT groups were compared using the Limma version of t-test. Proteins with a p-value < 0.05 calculated with limma were considered differentially expressed proteins and were used to perform enrichment analysis with IPA as for RNAseq data.

### Immunofluorescence

Briefly, neonatal hearts from 1-2 days pups were dissected in cold Hanks’ Balanced Salt Solution (HBSS) and the Pierce Primary Cardiomyocyte Isolation Kit (Thermo Fisher, 88281) was used to obtain neonatal cardiomyocytes. 2 x 10^6^ cells were seeded on p35 culture plates (Mattek, P35G-0-10-C) covered with matrigel solution (Fisher Scientific, 11553620). After two days in culture, cells were washed 3 times with PBS and fixed in 4% PFA for 10 min. Next, cells were permeabilized in 0.2% Triton X-100 in 1% BSA for 5 min and blocked with fetal bovine serum (FBS) for 1h at RT. Cells were incubated overnight at 4°C with mouse anti cMyBP-C monoclonal antibody (Santa Cruz Biotechnology, sc-137237, diluted 1:100 in FBS). After 3 washes with PBS, cells were incubated with chicken anti-mouse antibody conjugated to Alexa Fluor 647 (Life Technologies, Thermo Fisher, diluted 1:500 in FBS) for 1h at RT. Next, cells were washed three times with PBS and incubated with 5 µg/mL DAPI in FBS. Fluoromont G (bioNova, 0100-01) was used as mounting medium and allowed to cure overnight. Samples were imaged using Nikon A1-R confocal microscope with a Plan Apo100x/1.4 Oil objective. For image analysis, we used Fiji and a custom script written in imageJ macro language to automate creation and extraction of plot profile data from user defined lines, allowing the identification of all maximum and minimum intensity peaks. When calculating fluorescence intensity ratios, we considered the average value of the two maxima at the A band.

### Protein phosphorylation

Proteins were extracted from frozen left ventricle in a buffer containing 50 mM Tris-HCl pH 6.8, 10mM EDTA, 3% SDS, proteinase inhibitor cocktail set III (Millipore, 535140-1mL, diluted 1:100) and phosphatase inhibitor cocktail set II (Millipore, 524625-1set, diluted 1:100) using an automatic homogeniser (VWR, Pellet Mixer, 47747-370). Samples were left in ice for 30 min and then heated for 10 min at 60°C. Finally, they were centrifuged at 18,407 g for 5 min at RT to obtain supernatants. Delipidated and desalted samples were obtained from proteins extracts using a methanol-chloroform precipitation step. Protein concentration was determined using Bradford assay (Sigma, Cat. No. B6916). 10 μg from each sample was separated in 10% SDS-PAGE gels. After electrophoresis, gels were staining with Pro-Q Diamond (ThermoFisher, P33302) following manufacturer’s instruction. Next, gels were stained with Coomassie Coomassie Brilliant Blue R250 (Bio-Rad, Cat. No. 161-0436) and images were acquired using ChemiDoc system. Pro-Q intensity values of protein bands were normalized using Coomasie blue staining. Peppermint stick molecular weight marker (ThermoFisher, P27167) was used as a positive control for phosphorylation.

### Protein expression and purification

N-terminal His-tagged C3 and C-terminal His-tagged C0C7 domains of human WT and R502W cMyBP-C were produced and purified as described previously with some modifications ^11,24^. Briefly, Rosetta (DE3) *E.coli* carrying expression vectors pQE80L containing C3 WT or R502W ^23^ or pET 21a containing C0C7 WT ^11^ or R502W (introduced by site-directed mutagenesis by PCR) were grown to an absorbance at 600 nm of 0.6-1. Next, cells were induced with 1 mM IPTG for 3 h at 37 °C and at 250 rpm. For C3 domain purification, cells were lysed by sonication in 50 mM sodium phosphate pH 7, 300 mM NaCl, 1 mM DTT,10 µM leupeptin, 1 mM PMSF, 100 µg/mL lysozyme, 1% Triton X-100, 5 µg/mL DNAse I, 5 µg/mL RNAse A, 10 mM MgCl_2_. For C0C7 constructs the lysis buffer was 10 mM imidazole/50 mM NaH_2_PO_4_ pH 7.5, 300 mM NaCl, 1 mM β-mercaptoethanol, 0.01 mg/mL leupeptin, 1 mM PMSF and Roche complete protease inhibitor cocktail. C3 constructs were purified from the soluble fractions by nickel-based affinity (Ni-NTA, Qiagen). The most concentrated fraction was further purified by size-exclusion chromatography (SEC) on an FPLC using a Superdex 200 HiLoad 16/600 column in 10 mM HEPES, pH 7.2, 150 mM NaCl, 1 mM EDTA. C0C7 proteins were purified by one round in Ni-NTA column followed by ion-exchange Q-column and a final SEC on Superdex 200 HiLoad 16/600 in 10 mM imidazole pH 7.5, 100 mM potassium acetate, 4 mM MgCl_2_, 1 mM EDTA, 1 mM DTT.

Major peak fractions were pooled and used for MST. Purity of all samples before each purification step was evaluated using SDS-PAGE electrophoresis gels. Human β-cardiac myosin preparations were expressed using a modified AdEasy^TM^ Vector System (Qbiogene Inc) in differentiated C2C12 myoblasts as described elsewhere ^41^. In brief, two different truncated versions of myosin heavy chain were used: short S1 (sS1) consisting of residues 1 to 808 encoding one catalytic head, and 15-hep, which corresponds to residues 1 to 942 encompassing both S1 catalytic head and the first 15 heptad repeats of the proximal S2 coiled-coil region. Both constructs were co-expressed with human ventricular essential light chain 3. In the case of the 15-hep construct, there is an additional step that replaces mouse skeletal regulatory light chain by human, *E.coli*-expressed ventricular regulatory light chain.

### Microscale thermophoresis

For each interaction tested, cMyBP-C constructs were fluorescently labeled via His-tag using a commercially available kit (NanoTemper Technologies). Unlabeled sS1 or 15-hep myosin was titrated against 84 nM fluorescently labeled C3 or C0C7 protein. Sixteen serially diluted titrations of sS1 or 15-hep myosin were prepared to generate one full binding isotherm. All binding reactions were performed in 10 mM Tris-HCl pH 7.5, 4 mM MgCl_2_, 1 mM EDTA, 1 mM DTT, 50 mM potassium acetate, 0.1 mM sodium orthovanadate, 0.05% Tween-20. After mixture incubation for 30 min at 23 °C in the dark, samples were loaded into NT.115 premium-treated capillaries (NanoTemper Technologies) and then mounted in a Monolith NT.115 apparatus (Nanotemper Technologies) for binding measurements. To obtain binding isotherms, C3 or C0C7 fluorescence was measured in 5-8 replicate runs using a red LED (excitation 605–645 nm; emission 680–685 nm) at 40 – 80% excitation power and IR laser was used at 60% power. All data were acquired at 23 °C. Data analysis was carried out with NTAffinityAnalysis software (NanoTemper Technologies).

### Determination of myosin SRX/DRX ratios

To measure myosin SRX/DRX ratios, we followed published protocols ^14,16^. Briefly, ∼10 mg of left ventricular myocardium from cryopreserved hearts were cut and immersed in 10 mM potassium phosphate pH 7.0, 100 mM NaCl, 8 mM MgCl_2_, 5 mM EGTA, 3 mM NaN_3_, 5 mM ATP, 1 mM DTT, 20 mM 2,3-butanedione monoxime (BDM), 0.1% Triton X-100 (permeabilization buffer) on a rocker machine for 6 h at 4°C using fresh solution every 2 h. Then, the buffer was exchanged to 5 mM potassium phosphate pH 6.8, 120 mM potassium acetate, 5 mM magnesium acetate, 50 mM Mops, 5 mM ATP, 20mM BDM, 2 mM DTT, 50% (v/v) glycerol (glycerinating solution), and samples were equilibrated overnight in ice and used within two days. Small bundles of fibres were dissected manually in glycerinating solution at 4°C under a stereomicroscope. These skinned fibres were immobilized in a slide with double-sided adhesive tape and a flow cell was created with pre-cooled coverslips. Prior to recording, samples were further permeabilized for 30 min and washed in 5 mM potassium phosphate pH 6.8, 120 mM potassium acetate, 5 mM magnesium acetate, 50 mM Mops, and 2 mM DTT (rigor buffer). Next, permeabilized fibres were incubated with 250 μM mant-ATP in rigor buffer for at least 5 min before imaging. Fluorescence decay was recorded using a Nikon ECLIPSE Ti epifluorescence microscope coupled to an Orca ER hamamatsu CCD camera and a Plan Fluor 10x/0.3 Ph1 DLL objective. First, fibres were located using bright-field and then excited at 395 nm (DAPI filter) with 20 ms of exposure time. After imaging for 60 s, buffer exchange with 4 mM ATP in rigor buffer was done. Fluorescence decay was recorded during 10 min with 1 frame per second acquisition rate. For analysis, the average background intensity was subtracted from the fluorescence of regions of interest within muscle fibres. Resulting fluorescence decay curves were fit to the following double exponential function:

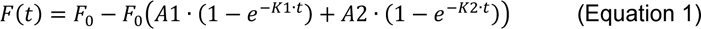

where 𝐹_O_ is the initial fluorescence, A1 and A2 are the weight of DRX and SRX states, and K1 and K2 are their rates of ATP turnover. Each individual experiment was fitted to this double exponential function using Igor Pro (Wavemetrics). To ensure convergence of the fitting procedure, we considered that 𝐾1 = 10 · 𝐾2 ^9^.

### Mavacamten treatment and plasma quantification

Mavacamten was acquired from MedChemExpress and was dissolved at 0.2 mg/mL concentration in 10% DMSO. Mice received 2.5 mg/kg/day of mavacamten or vehicle (H_2_O with 1% DMSO) in drinking water (daily replenishment of solutions). To quantify plasma levels of the drug by LC-MS, blood was collected by maxillary/facial vein puncture using a lancet in an EDTA containing tube. Plasma was obtained by centrifugation and stored at −80°C. 50-µL plasma aliquots were thawed in ice for 30 min and proteins were precipitated by adding a mixture of methanol:ethanol (1:1, v/v) at −20°C in a 5:1 ratio (solvent:sample). Samples were kept in ice for 20 min and centrifuged at 16,400 g for 20 min at 4°C. Supernatants were collected, transferred to a 1.5 mL tube and dried out in a speedvac (Savant SPD131DDA concentrator, Savant RVT5105 refrigerated vapor trap and OFP400 vacuum pump, ThermoFisher) for 2 h at RT. Finally, samples were resuspended in 50 µL acetonitrile:water (20:80, v:v) by constant shaking at 600 rpm for 10 min at 8°C and then centrifuged at 16,400 g for 5 min to remove insoluble material. Quantification of mavacamten was performed using an Ultimate 3000 HPLC system consisting of a degasser, two binary pumps, and thermostated autosampler, maintained at 8°C and coupled to an Orbitrap ELITE™ Hybrid Ion Trap-Orbitrap Mass Spectrometer (ThermoFisher Scientific). 5 µL of extract were injected into an Agilent Poroshell 120 EC-C18 column (2.1 x 100 mm, 4 µ) at 55°C. Mavacamten was eluted at 300 mL/min using (A) water with 0.1% formic acid and (B) acetonitrile with 0.1% formic acid. The gradient started from 5% to 95% of B in 20 min, keeping at 95% B for 6 min and returning to starting conditions in 0.2 min. Finally, re-equilibration at 5% of B was done for 8.8 min. Data were collected simultaneously in positive ESI full scan mode and PRM mode targeting m/z 274.1558 with 40 NCE at 60,000 resolution. Standard curve of mavacamten was analyzed alongside the plasma sample and peaks corresponding to mavacamten were integrated using FreeStyle 1.6 (ThermoFisher).

### Exercise capacity test

Exercise capacity was tested according to Dougherty *et al.* ^72^. Mice were placed on a treadmill with 5% inclination and an electric shock grid at the base. Prior to testing, mice were trained for 3 consecutive days with a full day of rest prior to exercise capacity evaluation. During each session, the treadmill speed was set at 20 cm/s initial speed and increased in steps to a final 40 cm/s speed (**Figure 7F**). Mice were removed from the treadmill if they spend more than 5 s uninterruptedly in the fatigue zone, which is defined as the part of the treadmill that includes 1 mouse body length from the shock grid as well as the grid itself. Exercise capacity was considered to be the total distance run by the mice up to the moment they were removed from the treadmill.

### Statistical analysis

Unless stated otherwise, data were expressed as mean ± SEM. Shapiro-Wilk (n ≤ 5) or Kolmogorov-Smirnov (n > 5) statistical tests were used to determine whether the data are normally distributed. Comparisons between 2 groups were done using parametric T-test when the data distribution was normal. Data not following normal distribution were analyzed by Mann-Whitney U-test. Two-way ANOVA with repeated measurements was used to compare distribution of data from wild-type and mutant groups across time. If repeated measurements were not available, a mixed model design was used. Statistical analyses were done using GraphPad Prism version 9.3.1. Comparisons with p-value < 0.05 were considered statistically significant.

### AUTHOR CONTRIBUTIONS

J.A.-C. conceived and led the project. A.F.-T and M.R.-P generated R502W mice. L.S-M, A.F.-T, M.A.L.-U., D.S.-O., J.A.N.-Á., M.S.-D, S.S., M.D., M.V.-O., G.G., E.C., R.B.-V, J.V., F.S.C., A.H., L.C. and J.A.-C designed mouse phenotyping and contextualized results. L.S.-M, A.F.-T, M.A.L.-U, A.F., V.L.-C, N.V., D.V.-C, L.S.-G, L.C., M.V.-O., and E.C. contributed to characterize the phenotype of mouse models. L.S.-M., D.P., J.A.S., and K.M.R. characterized protein binding by MST. L.S.-M and J.A.-C. drafted the manuscript with input from all authors. All authors agreed with the final version of the manuscript.

## ACKNOWLEDGMENTS

J.A.-C. acknowledges funding from the Ministerio de Ciencia e Innovación (MCIN, MCIN/AEI/10.13039/501100011033) through grants PID2020-120426GB-I00 and RED2022-134242-T, the Regional Government of Madrid (grant Tec4Bio S2018/NMT-4443, 50% co-financed by the European Social Fund and the European Regional Development Fund for the programming period 2014-2020), MCIN’s Severo Ochoa Program SEV-2015-0505 through a CNIC intramural grant 03-2016 IGP, and the European Research Council (ERC) under the European Union’s Horizon 2020 research and innovation programme (grant agreement No. [101002927]). J.V. acknowledges grants PID2021-122348NB-I00 from MICIU/AEI/ 10.13039/501100011033 and from “ERDF A way of making Europe”, PLEC2022-009298, PLEC2022-009235 and EQC2021-007053-P from MICIU/AEI/10.13039/501100011033 and from “European Union NextGenerationEU/ PRTR”, S2022/BMD-7333-CM (INMUNOVAR-CM) from the Regional Government of Madrid, and LCF/PR/HR22/52420019 from ”la Caixa” Foundation. The CNIC is supported by the Instituto de Salud Carlos III (ISCIII), the MCIN and the Pro CNIC Foundation and is a Severo Ochoa Center of Excellence (grant CEX2020-001041-S funded by MCIN). L.S-M is the recipient of an FPI predoctoral fellowship (PRE2021-097336 funded by MCIN). M.A.L.-U. is the recipient of a Juan de la Cierva – Formación postdoctoral grant (FJC 047055-I funded by MCIN). L.S-G is supported by grant PTA2020-019067-I funded by MICIU/AEI/10.13039/501100011033. We thank all members of our laboratories for discussion and ideas. We thank the expert support from the staff of CNIC’s Transgenesis, Genomics, Comparative Medicine and Microscopy units. We thank J.D. Hourcade, L.M. Criado, J.M. Fernández-Toro and M. Siguero (CNIC) for advice on CRISPR-Cas9 genetic engineering, and J. Ochala (U. of Copenhagen) for experimental advice on myosin SRX/DRX determination. We thank input from Víctor López, Josu Erquicia and María Padilla (Res@CNIC fellows). Microscopy was conducted at the CNIC Microscopy & Dynamic Imaging Unit. Biomedical Imaging was conducted at the Advanced Imaging Unit of the CNIC using ReDIB ICTS infrastructure TRIMA@CNIC, MCIN with the support of ISCIII (PT20/00044), co-financed by the European Union through the European Regional Development Fund (ERDF, “A way of doing Europe”).

