## Supplementary Information for "Broad therapeutic benefit of myosin inhibition in hypertrophic cardiomyopathy"

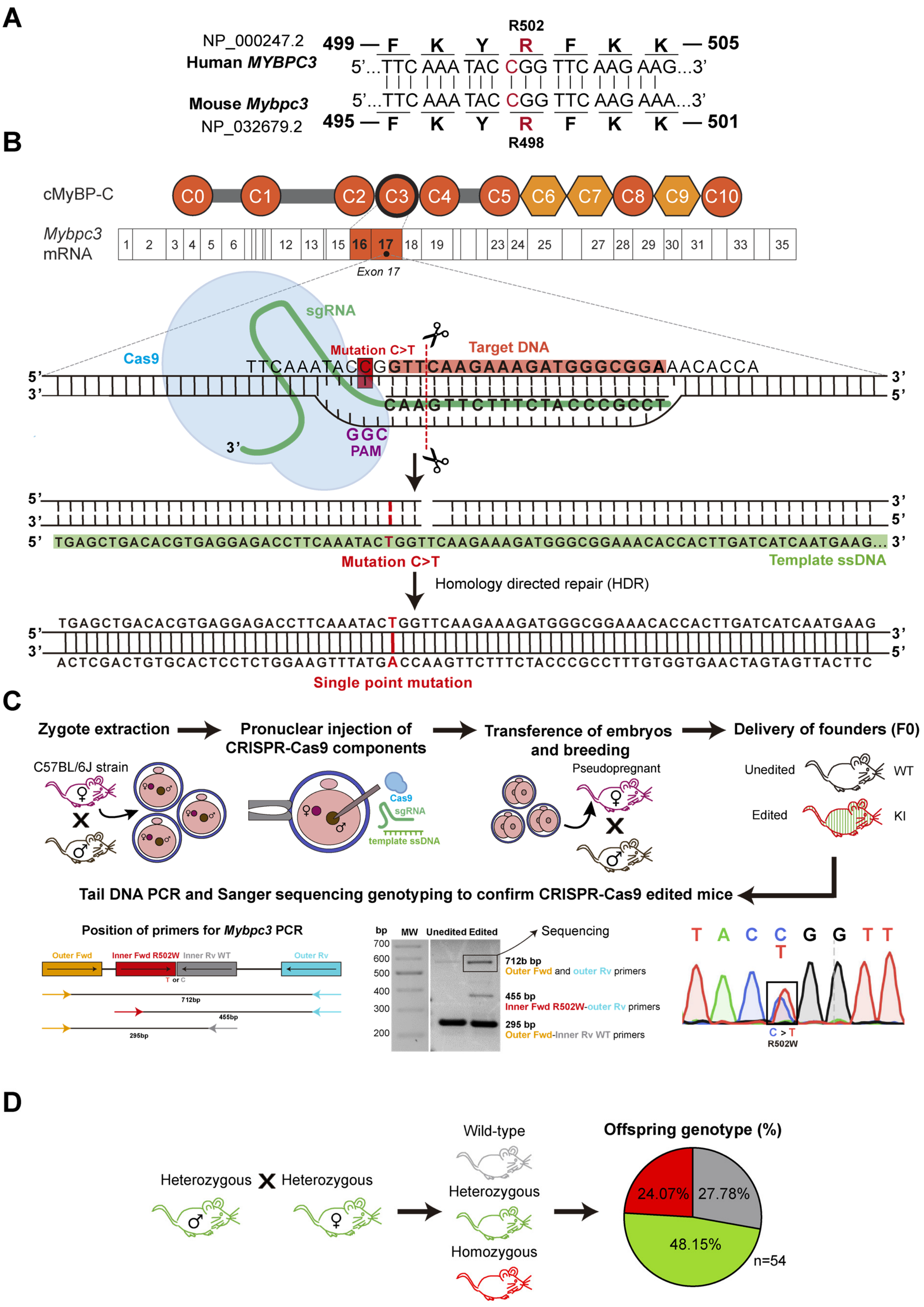


**Figure S1. Generation of R502W mice. A.** Nucleotide and protein sequence of the region targeted by cMyPB-C p.R502W variant. NCBI protein sequence entries are indicated. **B.** CRISPR-Cas9 strategy to insert the mutation R502W in cMyBP-C by homology directed repair. **C.** Scheme of the generation of mutant mice by pronuclear injection of CRISPR-Cas9 reagents and subsequent genotyping. **D.** Offspring genotype of R502W mice follows Mendelian proportions.

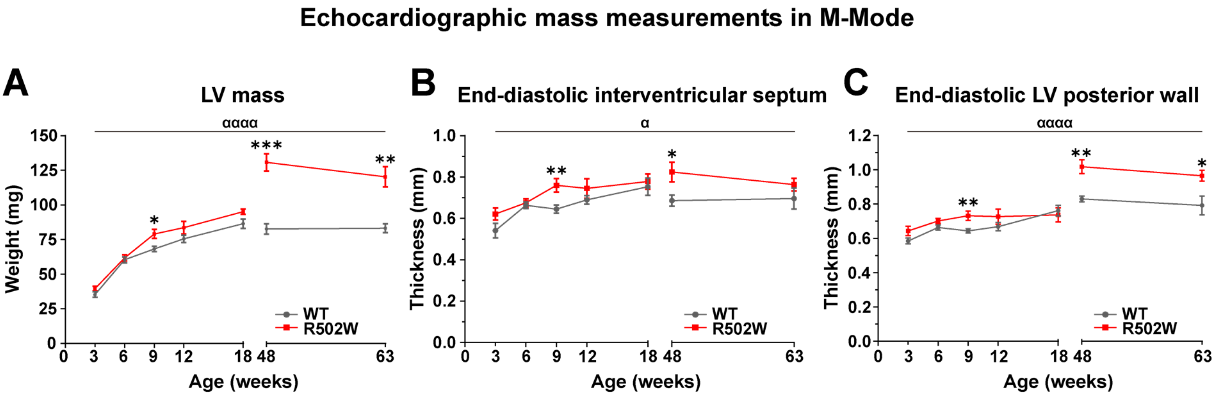


**Figure S2. Left ventricle anatomical parameters obtained in M-mode echocardiography for WT and R502W mice. A.** Left ventricle (LV) mass. **B.** End-diastolic interventricular septum thickness. **C.** End-diastolic LV posterior wall thickness. n = 5 – 20 per each genotype and age, * $p < 0.05$, ** $p < 0.01$, *** $p < 0.001$, $^{\alpha}p < 0.05$, $^{\alpha\alpha\alpha\alpha}p < 0.0001$.

### Additional parameters of cardiac function by echocardiography

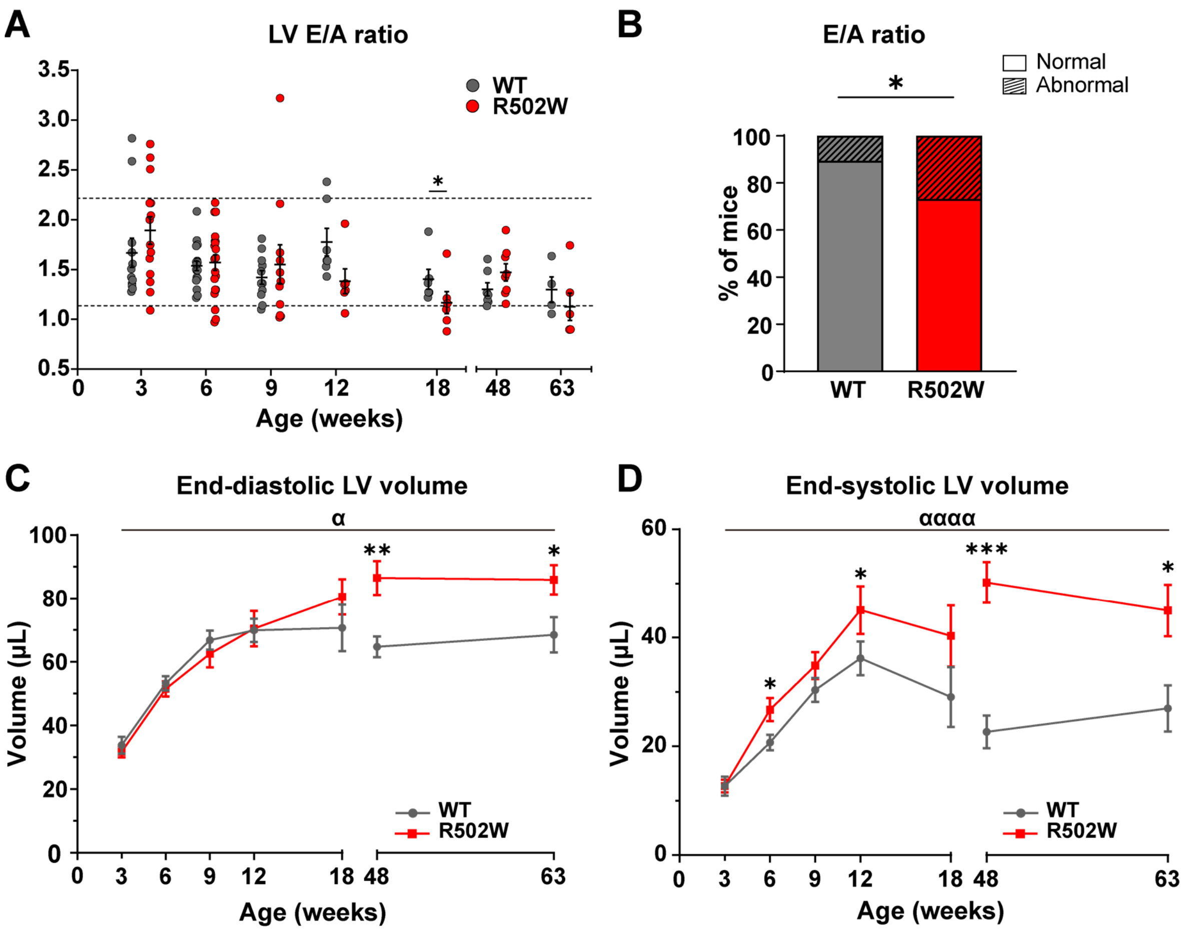


**Figure S3. Additional functional parameters in WT and R502W hearts by echocardiography. A.** Left ventricle (LV) E wave/A wave (E/A) ratio from pulsed-wave Doppler echocardiography. We chose as a normal range (indicated by dashed lines) data within the 5-95 percentiles of 6-to-12-week-old wild-type mice. **B.** Fraction of mice showing normal or abnormal E/A ratios (all ages, $n_{WT}$ = 66, $n_{R502W}$ = 71). Statistical significance assessed by Chi-square test. **C.** End-diastolic LV volume. **D.** End-systolic LV volume. Panels A, C and D: n = 5 – 20 per each genotype and age, * $p < 0.05$, **$p < 0.01$, ***$p < 0.001$, $^{\alpha}p < 0.05$, $^{\alpha\alpha\alpha\alpha}p < 0.0001$.

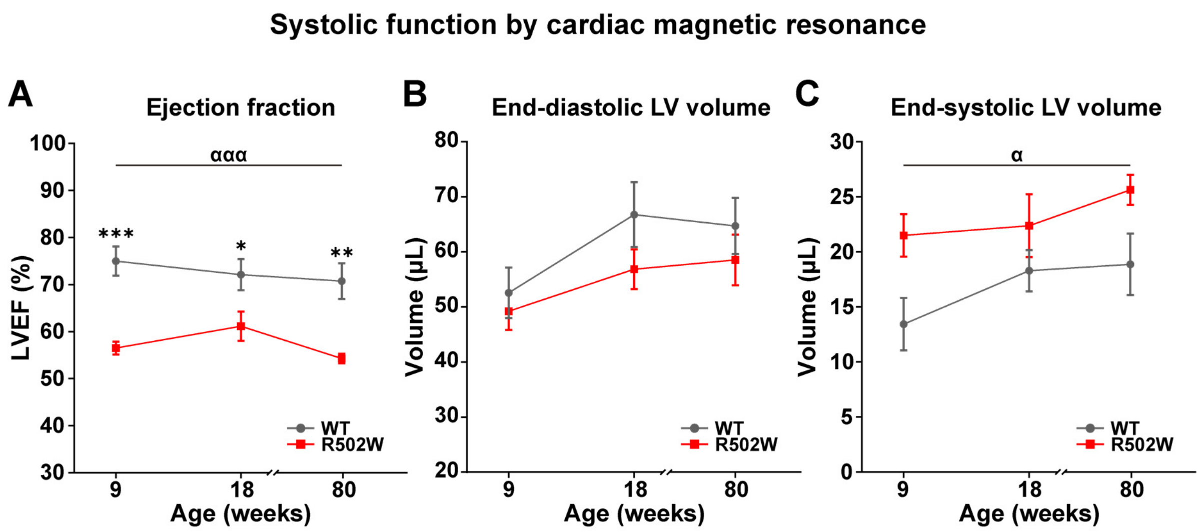


**Figure S4. Evaluation of systolic function using cardiac magnetic resonance. A.** Left ventricle ejection fraction (LVEF). **B.** End-diastolic LV volume. **C.** End-systolic LV volume. n = 5 per each genotype and age, $^{*}p < 0.05$, $^{**}p < 0.01$, $^{***}p < 0.001$, $^{\alpha}p < 0.05$, $^{\alpha\alpha\alpha}p < 0.001$.

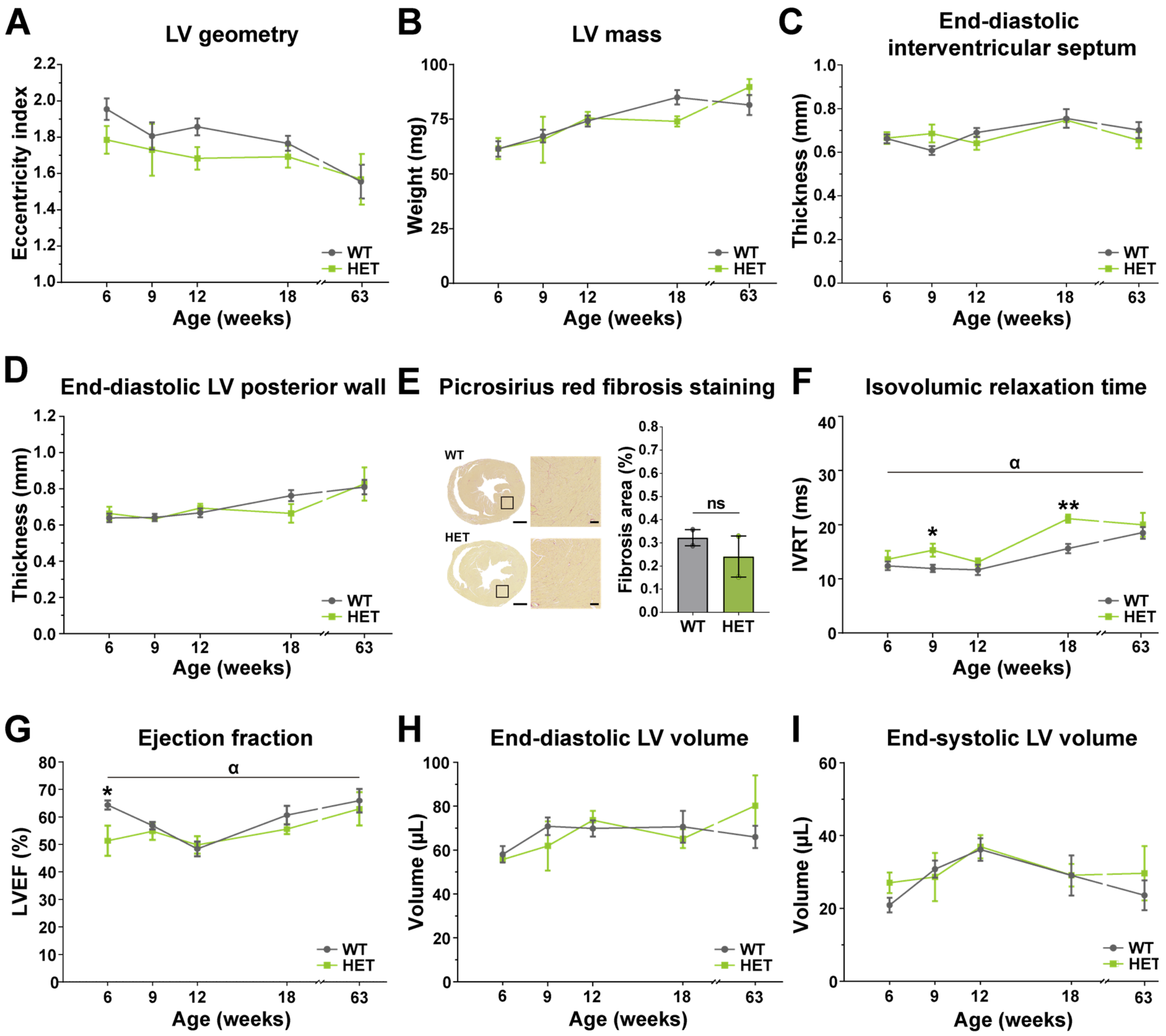


**Figure S5. Phenotyping of heterozygous R502W hearts by echocardiography and histology.** **A.** Left ventricle (LV) eccentricity index ratio. **B.** LV mass. **C.** End-diastolic interventricular septum thickness. **D.** End-diastolic LV posterior wall thickness. **E.** Percentage of fibrosis area by Picrosirius red staining of transversal cardiac paraffin sections (n= 2 animals per genotype). Scale bars are 1 mm (100 µm in insets). **F.** Isovolumic relaxation time. **G.** Left ventricle ejection fraction (LVEF). **H.** End-diastolic LV volume. **I.** End-systolic LV volume. n = 3 – 7 per each genotype and age for echocardiography measurements, * $p < 0.05$, ** $p < 0.01$, *** $p < 0.001$, $^{\alpha}p < 0.05$, ns: non-significant.

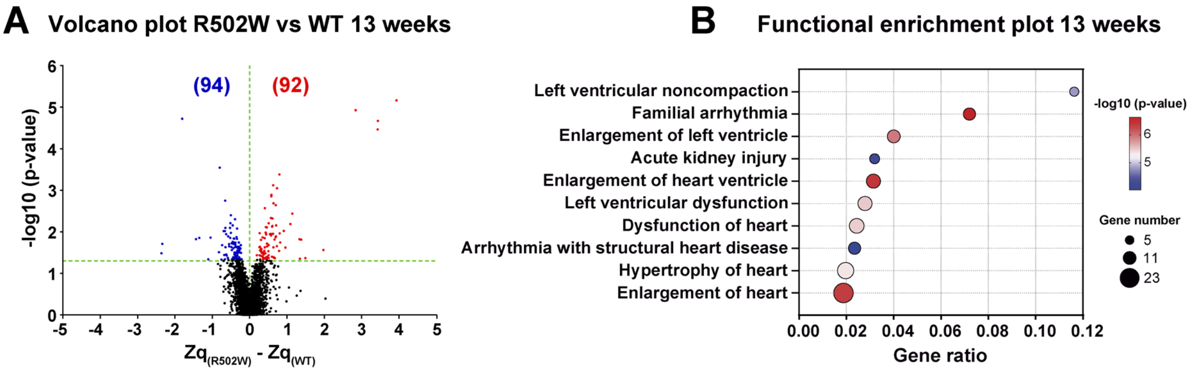


**Figure S6. Proteomics analysis of 13-week-old WT and R502W hearts. A.** Volcano plot showing differentially expressed proteins in R502W vs WT hearts. Upregulated proteins are shown in red and downregulated ones, in blue (n=3 per genotype). **B.** Functional enrichment bubble plot with the top 10 toxicology functions in Ingenuity Pathway Analysis.

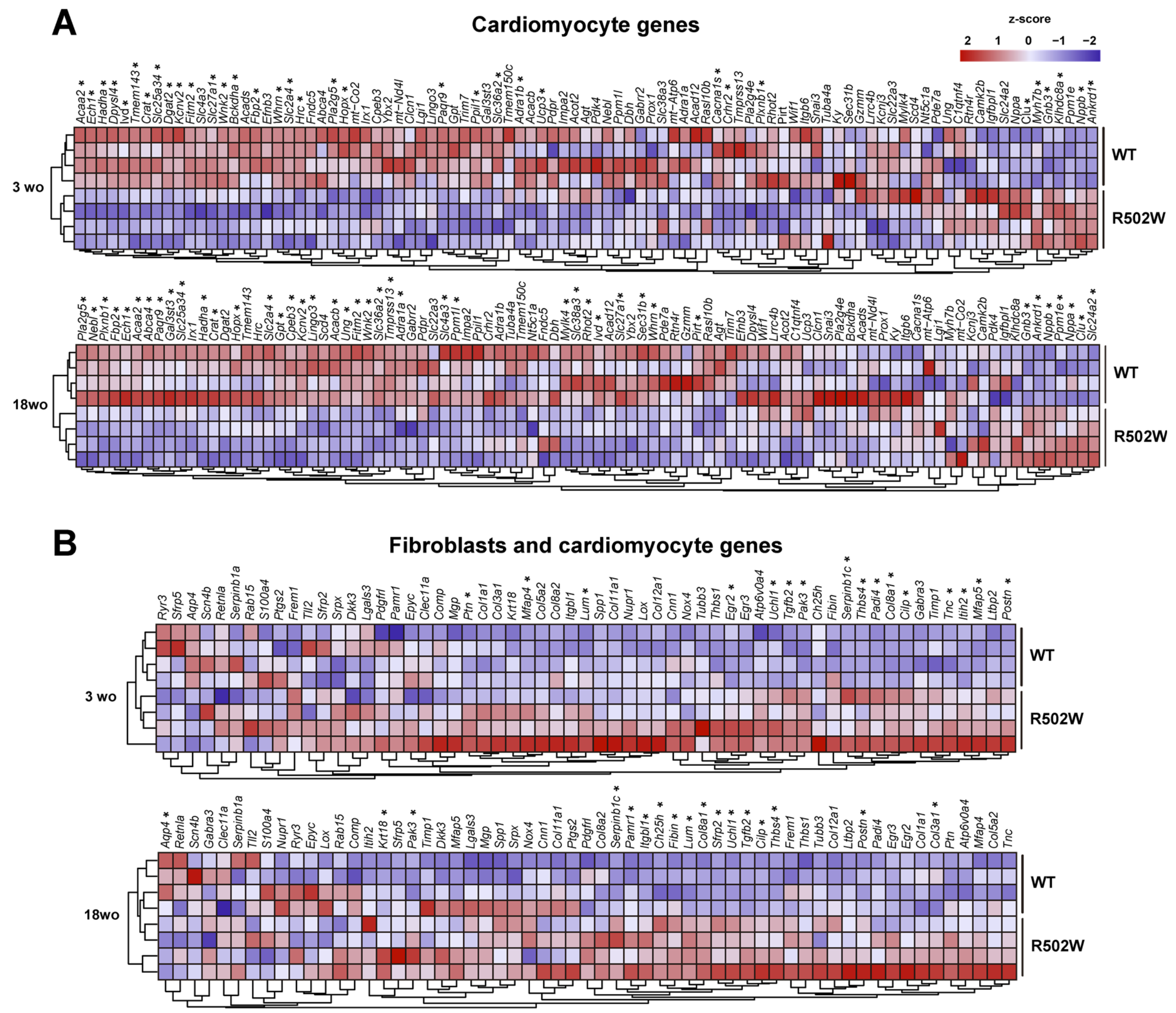


**Figure S7. Cluster analysis showing the expression in WT and R502W myocardium of the most dysregulated genes in HCM mouse models induced by mutations in myosin [1]. A.** Heatmaps depicting the expression levels of 93 cardiomyocyte genes in 3- and 18-week-old WT and R502W animals. **B.** Heatmaps depicting the expression levels of 59 genes found in fibroblasts or cardiomyocytes in 3- and 18-week-old WT and R502W animals. * Indicate genes with adjusted pvalue < 0.05 in R502W vs WT animals.

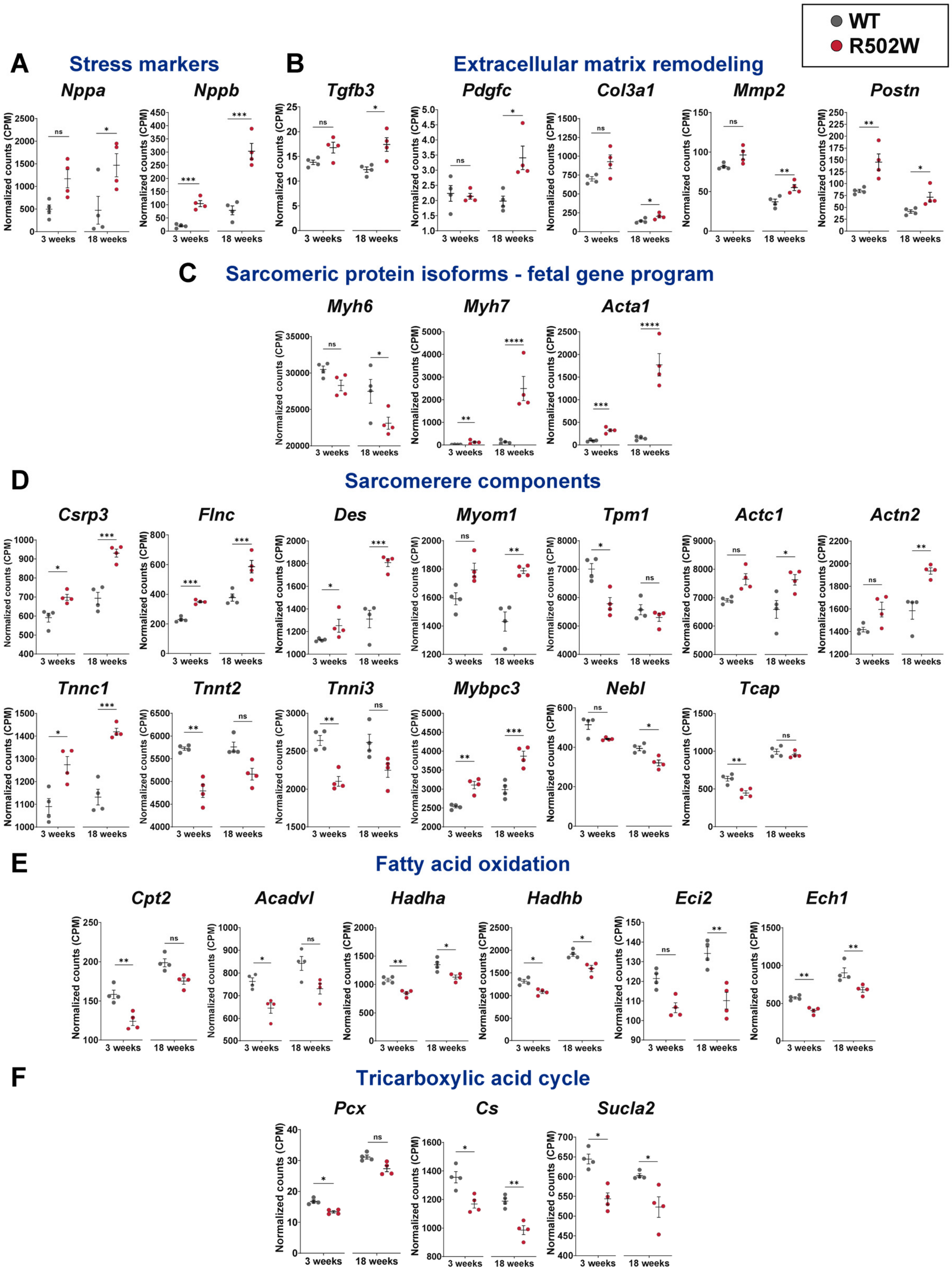


**Figure S8. Expression levels of genes involved in HCM-related pathways in 3- and 18-week-old WT and R502W mice (1/2). A.** Expression levels of stress markers typical of HCM. **B.** Expression levels of genes associated with extracellular matrix remodeling. **C.** Expression levels of genes coding for sarcomere proteins typical of the fetal gene transcriptional program. **D.** Expression levels of components of the sarcomere. **E.** Expression levels of genes belonging to the fatty acid oxidation pathway. **F.** Expression levels of genes implicated in the tricarboxylic acid cycle. *adjusted p-value < 0.05, **adjusted p-value < 0.01, ***adjusted p-value < 0.001, ****adjusted p-value < 0.0001, ns: non-significant.

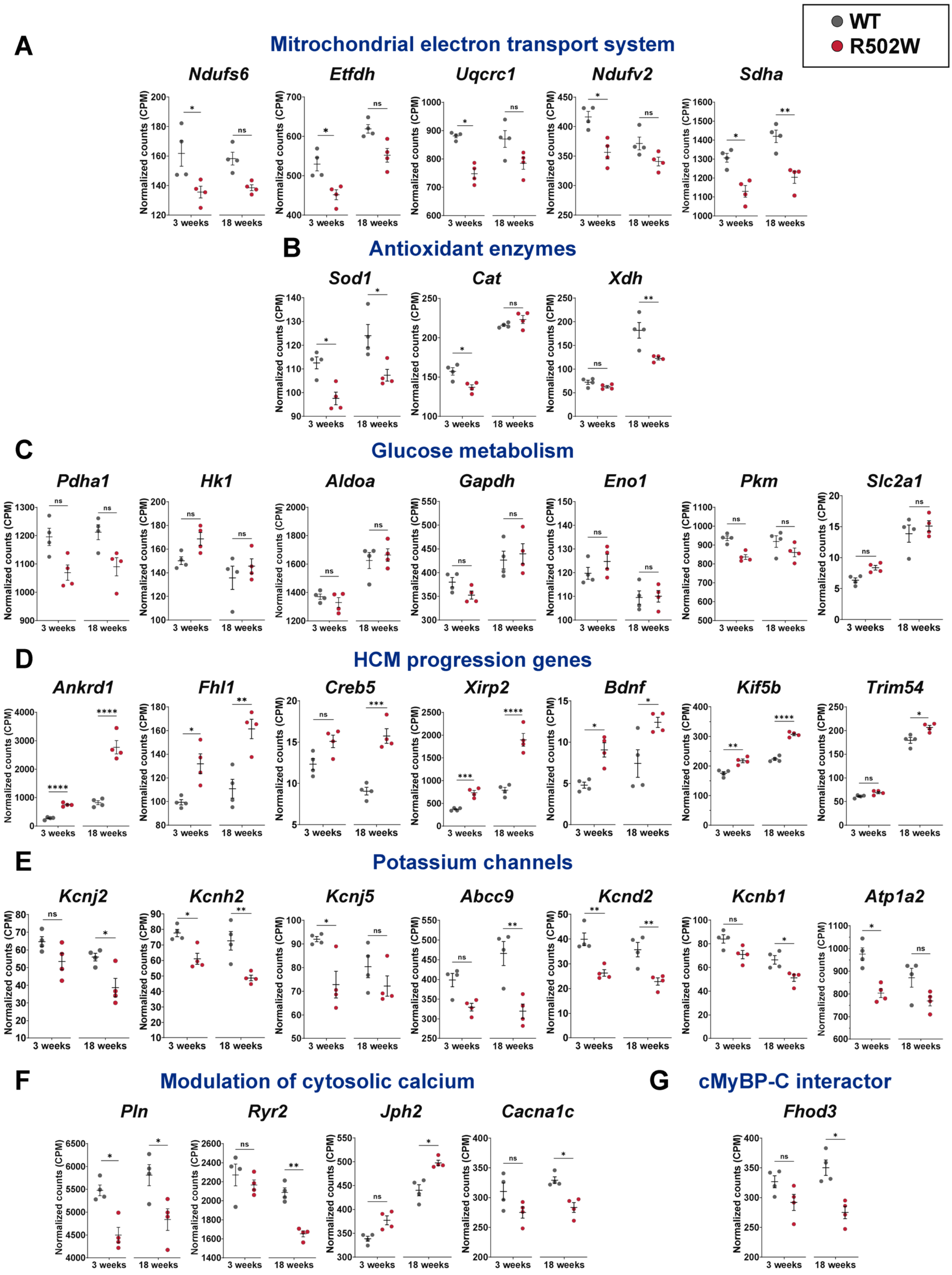


**Figure S9. Expression levels of genes involved in HCM-related pathways in 3- and 18-week-old WT and R502W mice (2/2). A.** Expression levels of genes associated with the mitochondrial electron transport system. **B.** Expression levels of genes encoding for antioxidant enzymes. **C.** Expression levels of genes relevant for glucose metabolism. **D.** Expression levels of other genes implicated in the progression of HCM. **E.** Expression levels of genes encoding for potassium channels. **F.** Expression levels of genes implicated in cytosolic calcium modulation. **G.** Expression level of a recently described interactor for cMyBP-C. *adjusted p-value < 0.05, **adjusted p-value < 0.01, ***adjusted p-value < 0.001, ****adjusted p-value < 0.0001, ns: non-significant.

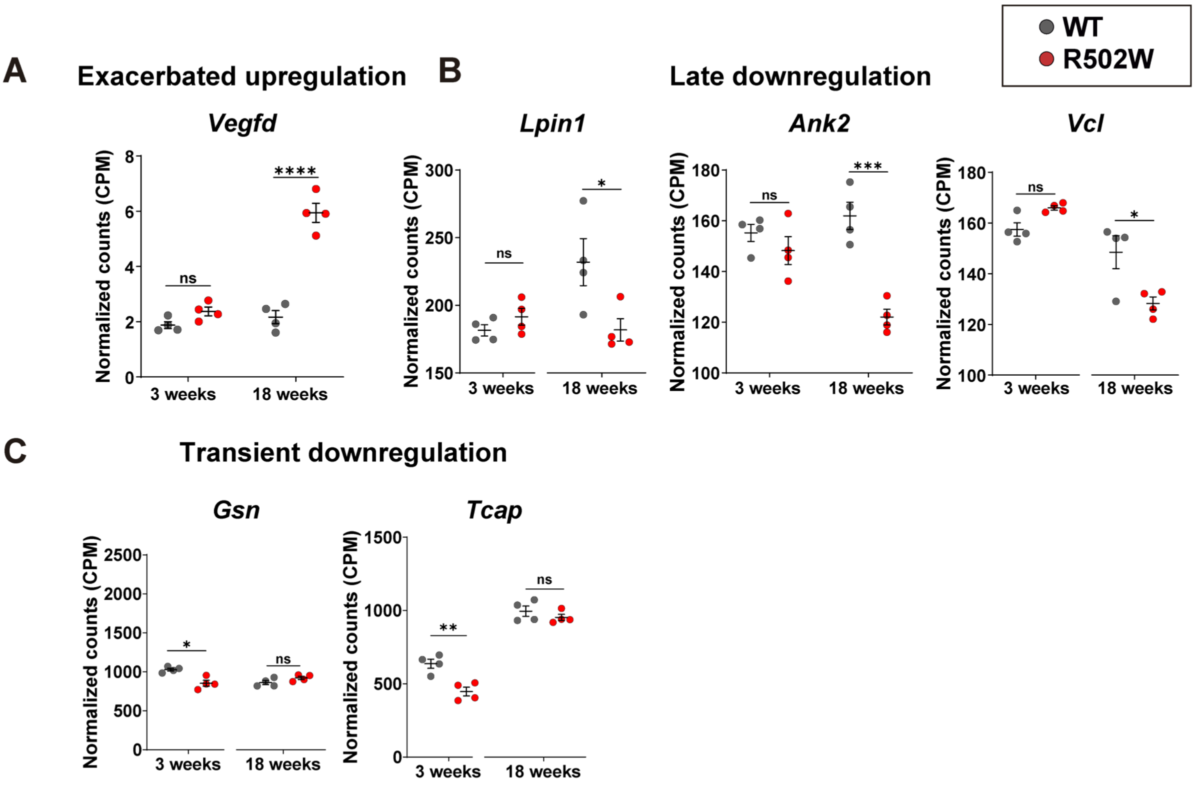


**Figure S10. Levels of selected HCM genes that show differential myocardial expression with respect to WT in 3- and 18-week old R502W animals. A.** Expression levels of *Vegfd*, showing exacerbated upregulation in 18-week-old R502W animals. **B.** Expression levels of three genes showing late downregulation in 18-week-old R502W animals. **C.** Expression levels of genes with a transient downregulation in 3-week-old R502W animals. *adjusted p-value < 0.05, **adjusted p-value < 0.01, ***adjusted p-value < 0.001, ****adjusted p-value < 0.0001, ns: non-significant.

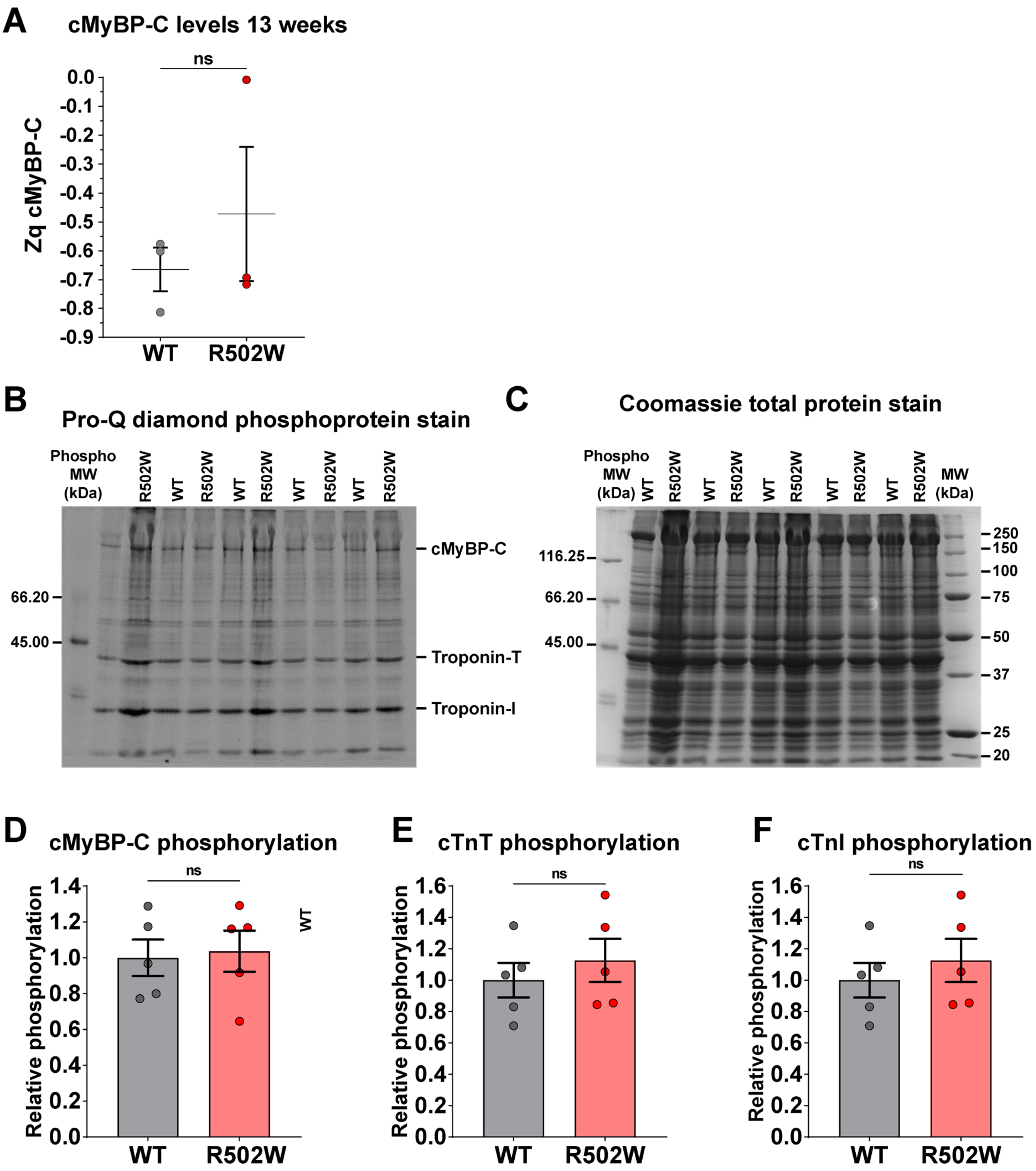


**Figure S11. Levels of cMyBP-C by mass spectrometry and phosphorylation of sarcomeric proteins. A.** cMyBP-C protein levels in 13-week-old WT and R502W mice from proteomics data (n = 3 animals per genotype). **B.** SDS-PAGE gel stained with ProQ diamond showing phosphorylation levels of protein extracts from 18-week-old WT and R502W animals. **C.** SDS-PAGE gel stained with Coomassie blue. **D.** Densitometry ratio of ProQ diamond/Coomassie blue staining to quantify phosphorylation levels of cMyBP-C. **E.** Densitometry ratio of ProQ diamond/Coomassie blue staining gels to quantify phosphorylation levels of troponin-T (TnT). **F.** Densitometry ratio of ProQ diamond/Coomassie blue staining gels to quantify phosphorylation levels troponin-I (TnI) [2]. Data in panels D-F correspond to n=5 animal per genotype and are referred to phosphorylation levels in WT mice. ns = non-significant.

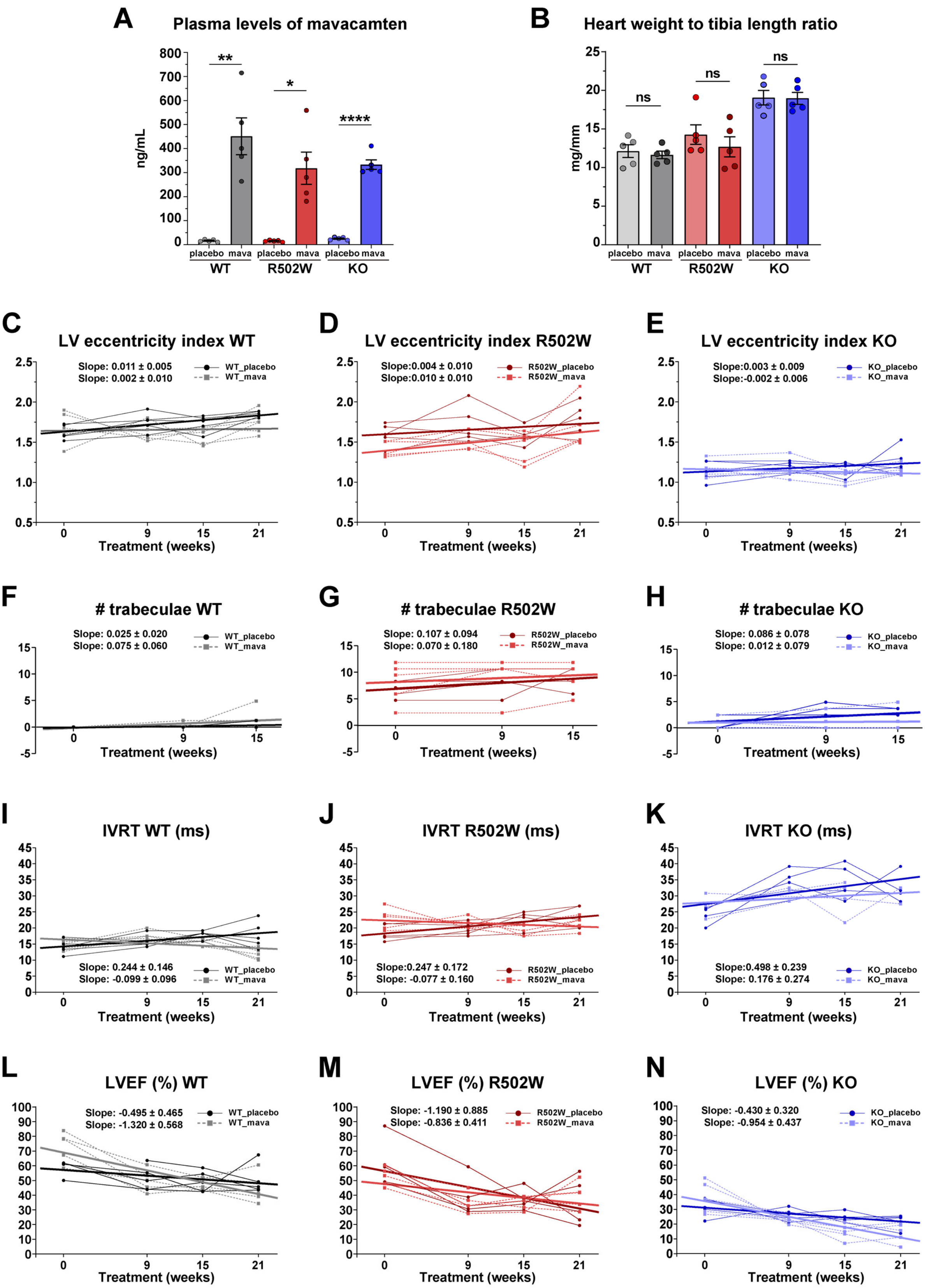


**Figure S12. Additional results from the experiment to test mavacamten efficacy. A.** Levels of mavacamten in plasma after four weeks of treatment. **B.** Heart weight to tibia length ratio. **C-E.**Left-ventricle eccentricity index ratio. **F-H.** Number of trabeculae evaluated by cardiac magnetic resonance **I-K.** Isovolumic relaxation time (IVRT) measured by echocardiography. **L-N.** Left ventricle ejection fraction (LVEF). n= 5 animals per time and group. In panels C-N, insets show the slopes of linear fits to the data with 83% confidence intervals. ****p < 0.0001, **p < 0.01, *p < 0.05, ns = non-significant.

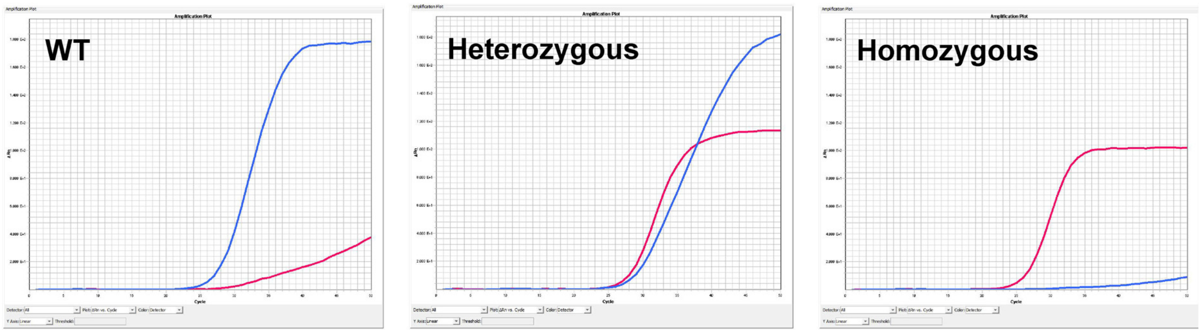


**Figure S13. Representative qPCR results for phenotyping R502W mice using rhAmp™ system.**

**Supplementary Table S1. Primers and probes used in this study.**

| | |
|---|---|
| sgRNA | 5’-TCCGCCCATCTTTCTTGAAC-3’ |
| Template ssDNA | 5’ TGAGCTGACACGTGAGGAGACCTTCAAATACTGGTTCAAGAAAGA TGGGCGGAAACACCACTTGATCATCAATGAAGCAACCCTGGAGGATGC AGGACACTATGCAGTACGCACAAGTGGAGGCCAG 3’ |
| Outer Forward cMyBP-C | 5’ AACTCAGGTTCAGGGGTTGAC 3’ |
| Outer Reverse cMyBP-C | 5’ CATCTTCGAGTCCGTCGGTG 3’ |
| Inner Forward R502W cMyBP-C | 5’ GCCCATCTTTCTTGAACCA 3’ |
| Inner Reverse WT cMyBP-C | 5’ TGAGGAGACCTTCAAATACC 3’ |
| rhAmp SNP allele specific WT primer | 5'-TGAGGAGACCTTCAAATACC-3' |
| rhAmp SNP allele specific R502W primer | 5'-GTGAGGAGACCTTCAAATACT-3' |
| rhAmp SNP locus specific primer | 5'-GCCGTACTGCATAGTGTCCTG-3' |
| Forward WT for KO cMyBP-C | 5-GAGATCCATGGAGGAGTGGA-3' |
| Reverse WT for KO cMyBP-C | 5'-TACCTAACCCTGCCCACAAG-3' |
| Forward KO for KO cMyBP-C | 5'-AGCCTTCTCTTCCAGCCCCAG-3' |
| Reverse KO for KO cMyBP-C | 5'-GCATCGCCTTCTATCGCCTTCTTGACG-3' |
